# Exploratory polarization facilitates mating partner selection in *Saccharomyces cerevisiae*

**DOI:** 10.1101/2020.09.07.285965

**Authors:** Manuella R. Clark-Cotton, Nicholas T. Henderson, Michael Pablo, Debraj Ghose, Timothy C. Elston, Daniel J. Lew

## Abstract

Yeast decode pheromone gradients to locate mating partners, providing a model for chemotropism. How yeast polarize toward a single partner in crowded environments is unclear. Initially, cells often polarize in unproductive directions, but then they relocate the polarity site until two partners’ polarity sites align, whereupon the cells “commit” to each other by stabilizing polarity to promote fusion. Here we address the role of the early mobile polarity sites. We found that commitment by either partner failed if just one partner was defective in generating, orienting, or stabilizing its mobile polarity sites. Mobile polarity sites were enriched for pheromone receptors and G proteins, and we suggest that such sites engage in an exploratory search of the local pheromone landscape, stabilizing only when they detect elevated pheromone levels. Mobile polarity sites were also enriched for pheromone secretion factors, and simulations suggest that only focal secretion at polarity sites would produce high pheromone concentrations at the partner’s polarity site, triggering commitment.

## INTRODUCTION

Directed growth (chemotropism) or movement (chemotaxis) in response to a chemical signal is critical for biological processes including aggregation in *Dictyostelium discoideum* (Nichols et al., 2015), pollen tube growth during plant fertilization (Higashiyama and Takeuchi, 2015), axon guidance during neural development (Bellon and Mann, 2018), and neutrophil migration in the mammalian immune response (Sarris and Sixt, 2015). However, the mechanisms by which cells choose the direction of polarized growth or movement are incompletely understood.

*Saccharomyces cerevisiae* exhibits polarized growth during mating, culminating in the fusion of two cells to form a zygote (Merlini et al., 2013). Because yeast are immotile, two cells must grow toward each other to facilitate membrane fusion. Cells of each mating type, **a** and α, secrete pheromones that are sensed by cognate G protein-coupled receptors on cells of the opposite mating type. Pheromone sensing triggers the activation of a MAPK cascade, cell cycle arrest in G1, increased transcription of mating-specific genes, and polarized growth toward the mating partner (Dohlman and Thorner, 2001).

Polarized growth is directed by the Rho-GTPase Cdc42, its regulators, and its effectors, which become concentrated at a small region of the cell’s cortex (Park and Bi, 2007) to form a “polarity site.” Polarity sites are assembled by a positive feedback mechanism in which active Cdc42-GTP binds the scaffold protein Bem1, promoting the activation of nearby inactive Cdc42-GDP to form an initial small cluster of polarity factors (Johnson et al., 2011; Kozubowski et al., 2008). Formins (effectors of Cdc42) trigger the orientation of actin cables toward the site, promoting vesicle-mediated delivery of cell wall-remodeling enzymes that enable polarized growth (Pruyne et al., 2004).

Yeast cells prefer partners that make more pheromone (Jackson and Hartwell, 1990a; Jackson and Hartwell, 1990b), and cells exposed to an exogenous gradient of pheromone tend to grow up-gradient (Segall, 1993). Consequently, models of partner selection often depict a cell polarizing growth up a gradient of pheromone that emanates from a single source (Arkowitz, 1999; Ismael and Stone, 2017) (***Fig. 1A***). However, a stable unidirectional pheromone gradient may be infrequent during mating in the wild. Meiosis of a diploid cell produces four haploid spores (two **a**, two α) encased in an ascus. Upon exposure to nutrients, germinating spores from a single ascus can mate with each other, but since each spore is adjacent to two potential partners (Taxis et al., 2005), it is unclear how the resulting pheromone landscape yields orientation toward one partner (Jin et al., 2011; Rappaport and Barkai, 2012) (***Fig. 1B***). Moreover, germinating spores derived from wild yeast strains often proliferate to form microcolonies (McClure et al., 2018) and switch mating type (Haber, 2012) before mating, so that each cell is likely to have more than one potential partner (***Fig. 1C***). Even when surrounded by many potential partners, cells choose only one (Jackson and Hartwell, 1990a; Jackson and Hartwell, 1990b). With multiple local pheromone sources that change as cells bud or mate, how do cells reliably orient polarity toward one partner?

**Fig. 1.**
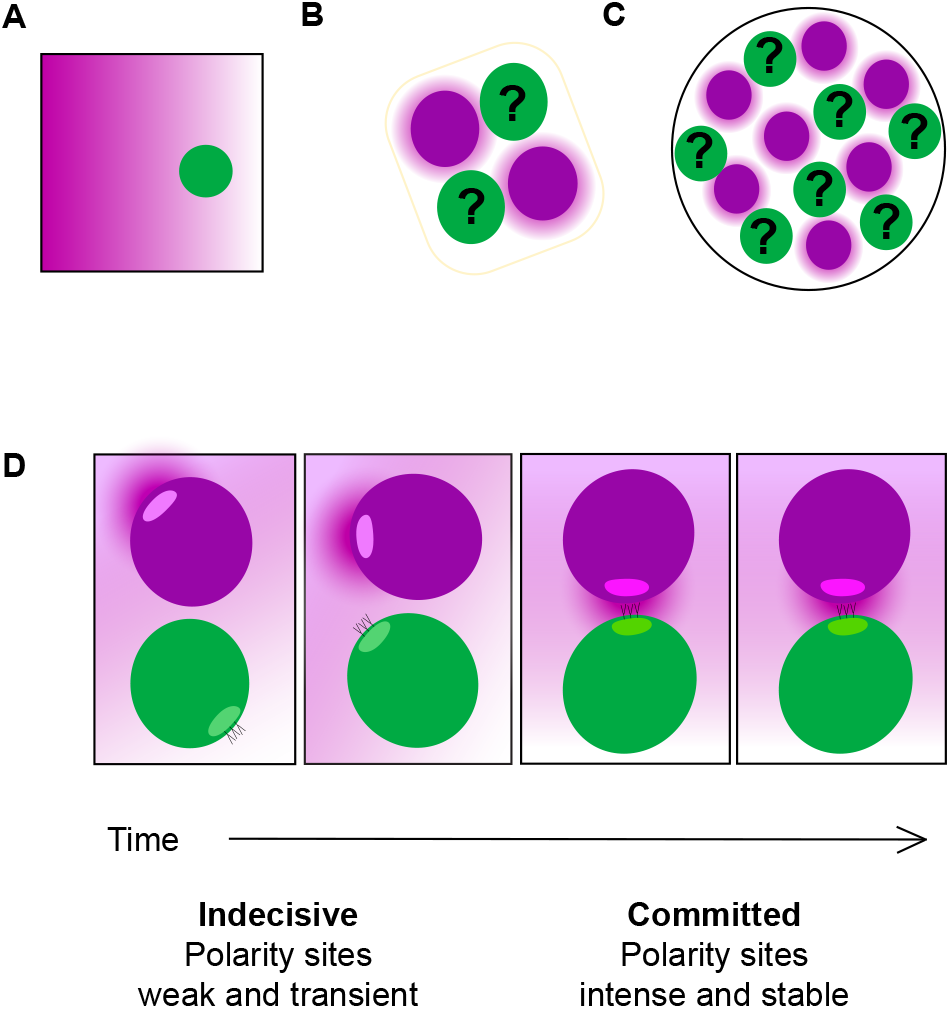
Pheromone landscapes encountered by yeast cells. (A) Stable unidirectional pheromone gradient, as generated by micropipet or microfluidics device. (B) Germinating spores in an ascus, where two potential partners (magenta) are expected to generate similar α-factor gradients, making them equally attractive to the **a**-cells (green). (C) Microcolony containing a mixture of **a**-(green) and α-cells (magenta). The proximity of multiple potential partners complicates the task of orienting toward a single partner. (D) Exploratory polarization model of partner selection. During the indecisive period (frames 1 and 2), diffusion of pheromone released at the α-cell’s (magenta) polarity site yields a low pheromone concentration at the **a**-cell’s (green) polarity site. When the two polarity sites are apposed, the a-cell senses a high concentration of pheromone. Both cells sense and secrete pheromone, but for simplicity, only the **a**-cell’s receptors and α-cell’s pheromone are shown.

For successful mating, the pheromone landscape must be decoded to orient polarity toward the partner. However, imaging of polarity factors revealed that initial polarity sites were not always oriented toward the eventual mating partner in mating mixes (Henderson et al., 2019; Wang et al., 2019) or up the gradient in cells exposed to artificial pheromone gradients (Dyer et al., 2013; Hegemann et al., 2015; Jin et al., 2011; Kelley et al., 2015; Vasen et al., 2020). Rather, the location of the polarity site changed over time, stabilizing at better-oriented locations. In mating mixes, weak clusters of polarity factors appeared, disappeared, and changed position in a chaotic manner (Henderson et al., 2019). After this “indecisive phase” of 10-120 min, cells developed strong, stable polarity sites oriented toward the partner, suggesting that they made a “commitment” to the partner. Similarly, cells exposed to a steep pheromone gradient in a microfluidics device spent a variable interval with weak and mobile polarity sites before developing a strong polarity site at a stable position (Hegemann et al., 2015). These studies suggested that yeast cells process spatial information about the local pheromone landscape during a search period of variable duration, then commit to a specific orientation for polarized growth.

Spatial information about the pheromone landscape could be extracted by “global” or “local” sensing strategies (Hegemann and Peter, 2017; Kelley et al., 2015; Martin, 2019). In global sensing, cells compare the concentration of ligand-bound receptors around the cell surface to infer the direction of the pheromone source. In local sensing, cells primarily detect pheromone in a sensitized zone centered around the polarity site, moving the site around to infer the direction of the pheromone source. These models are not mutually exclusive.

Evidence for global sensing comes from the observation that in mating mixes, initial weak polarity sites were oriented toward their eventual mating partners more often than would be expected by chance (Henderson et al., 2019). Thus, some spatial information was available before any polarity sites were visible. Other studies reported that initial polarity sites were frequently oriented toward bud site-selection cues rather than pheromone sources (Vasen et al., 2020; Wang et al., 2019). Regardless of initial orientation, global sensing during the indecisive phase (when polarity sites are weak and mobile) could promote the selection of optimal locations for stable “committed” polarity sites.

Evidence for local sensing comes from the observation that when a strong polarity site is present, pheromone receptors and associated G proteins accumulate around the polarity site (Ayscough and Drubin, 1998; McClure et al., 2015; Suchkov et al., 2010). Thus, cells with a strong polarity site sense pheromone preferentially in the vicinity of the site. It is unclear whether the weak and transient polarity sites characteristic of the indecisive phase would similarly enable local sensing.

The observations discussed above suggested a potential strategy for partner search that we call “exploratory polarization” (Henderson et al., 2019). One component of this hypothesis is local sensing: this proposes that cells sense pheromone primarily near the indecisive polarity sites. The second component is local pheromone secretion: this proposes that most pheromone is secreted from the indecisive polarity sites, so the pheromone landscape changes as the polarity sites move. When polarity sites in potential partners are not properly aligned, the pheromone released from one cell is dissipated by diffusion and detected at a low concentration at the partner’s polarity site (***Fig. 1D***). However, when two polarity sites face each other (and *only* in that case), a high concentration of pheromone is detected at each site (***Fig. 1D***). This stabilizes both sites, leading to spatially and temporally coordinated commitment by both partners. According to this hypothesis, successful communication between two cells requires the simultaneous coorientation of their polarity sites.

A similar strategy was proposed for *Schizosaccharomyces pombe*, where weak polarity sites formed and disappeared at different locations before commitment (Bendezu and Martin, 2013). Pheromone secretion and sensing factors were enriched at these transient sites, and computational models confirmed that local secretion and sensing/signaling would promote effective partner location in realistic geometries and timescales (Merlini et al., 2016). Thus, an exploratory polarization strategy might be conserved between evolutionarily distant fungi (Martin, 2019).

In this study, we tested the ability of wildtype cells to locate partners that are impaired in either the formation, localization, or stabilization of weak, indecisive phase polarity sites. Wildtype cells appeared to search for an extended indecisive period but did not commit (polarize stably) to the aberrant partners. We also confirmed that transient polarity sites are enriched in pheromone sensing, signaling, and secretion proteins. Simulations of the pheromone landscape based on documented pheromone secretion rates provide quantitative support for the idea that pheromone levels sufficient to promote commitment are only achieved when polarity sites of mating partners become aligned. We conclude that local sensing and secretion by two cooriented polarity sites enables commitment by a mating pair.

## RESULTS

### Behavior of mating cells before commitment to a partner

Reports of polarity before commitment vary, so we began by re-examining this behavior. Imaging of polarity probes like Bem1 (***Fig. 2A***) in wildtype cells showed weak initial polarity sites that exhibited chaotic “indecisive” behavior before becoming strong, stable polarity sites facing the partner (Henderson et al., 2019). However, imaging of probes for sensing/signaling proteins (***Fig. 2A***) suggested that cells undergo a more stereotyped behavior, with initial polarization always adjacent to the previous site of cytokinesis, followed by steady unidirectional tracking toward the eventual mating partner (Wang et al., 2019). We considered the possibility that probe selection might affect conclusions about cells’ behavior prior to commitment.

**Fig. 2.**
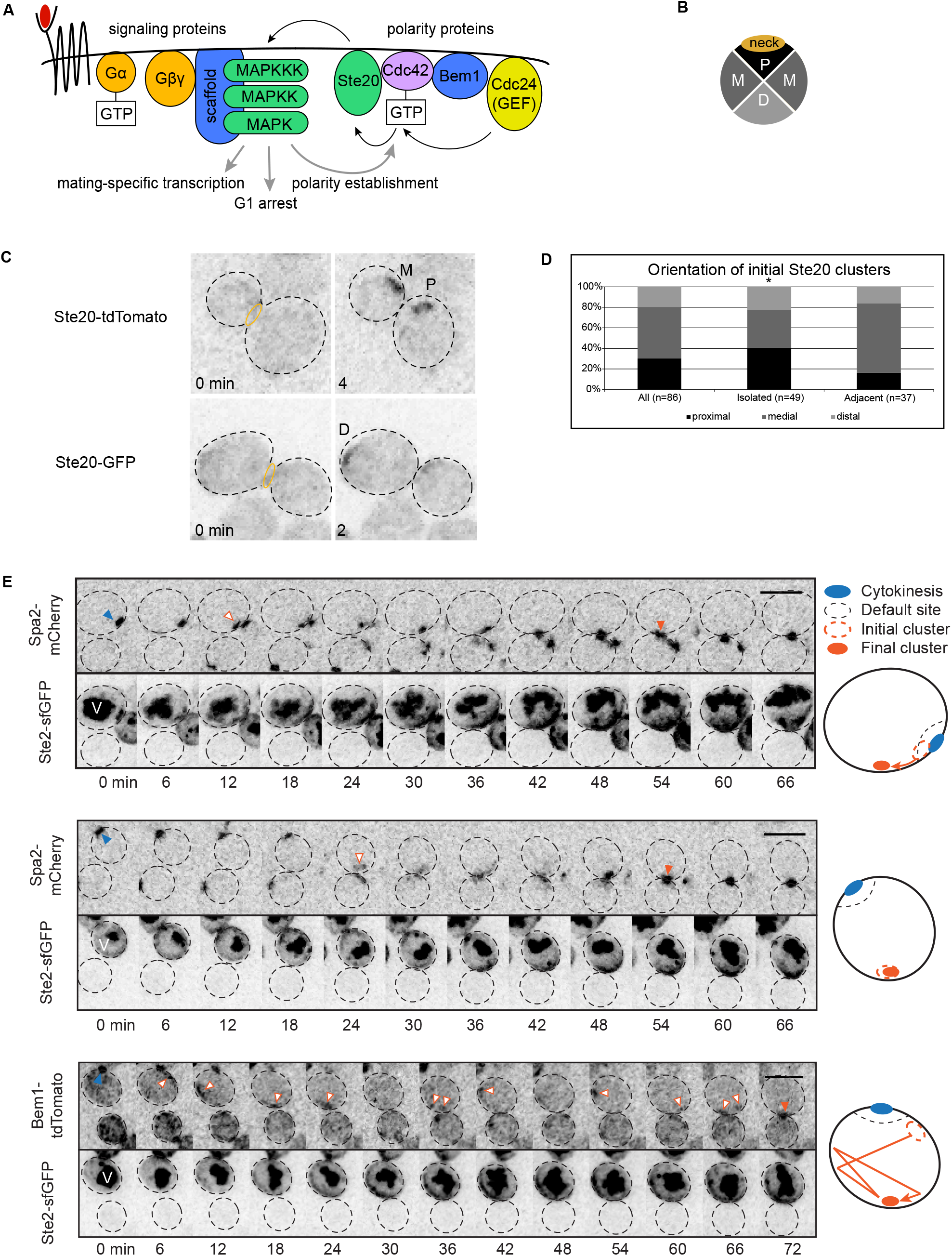
Behavior of mating cells before commitment to a partner. (A) Selected signaling and polarity proteins. When pheromone (red) binds receptor, free Gβγ recruits the scaffold to the membrane, activating the MAPK to promote downstream events, including polarization. (B) Cartoon depicting the quadrants (proximal, medial, distal) used to score the initial orientation of Ste20 relative to the mother-bud neck. If Ste20 polarizes randomly, then we expect 25% proximal, 25% distal, and 50% medial. (C) MAT**a***STE20-GFP* (DLY14364) and MATα *STE20-tdTomato* (DLY14413) cells were mixed and imaged during mating. Inverted maximum-projection montages illustrate representative mother-daughter pairs. *Top*, Ste20-tdTomato formed clusters at medial (M) and proximal (P) sites. *Bottom*, a Ste20-GFP cluster formed distal (D) to the neck (yellow oval). (D) Initial orientation of Ste20 clusters from the mating mix in (C), and in subsets that were (adjacent) or were not (isolated) touching potential mating partners. *: goodness-of-fit test, p = 0.05. (E) Cells harboring *STE2-sfGFP* and either *BEM1-tdTomato* (DLY22243) or *SPA2-mCherry* (DLY20712) were mixed with cells of the opposite mating type (*BEM1-tdTomato*, DLY22340 or *SPA2-mCherry*, DLY8503, respectively) and imaged during mating. Following Ste2 degradation, sfGFP accumulates in the vacuole (V). Ste2-sfGFP crescents gradually intensified in regions visited by polarity clusters. Blue arrowheads, accumulation of Spa2 or Bem1 at cytokinesis site; white arrowheads, polarity sites during indecisive phase; orange arrowheads, final polarity site. Cartoons summarize polarity behaviors. Scale bars: 5 μm.

The Bem1 probe concentrates at the mother-bud neck during cytokinesis. To control for possible bias against neck-proximal initial polarization, we visualized the Cdc42 effector Ste20, which does not concentrate at the neck during cytokinesis. Scoring of initial Ste20 polarization did not reveal a significant bias to any sector (***Fig. 2B-D***) (p = 0.40, n = 86). Analyzing cells that were or were not directly adjacent to a potential partner (“isolated” vs. “adjacent”, ***Fig. 2C,D***) revealed that isolated cells showed a non-random distribution (p = 0.05, n = 49), while adjacent cells did not (p = 0.14, n = 37). Thus, under our mating conditions, isolated cells might have a mild preference for the default site, but those that are near a mating partner do not.

Following initial polarization, Ste20 displayed indecisive behavior similar to that of Bem1 (**Movie 1**), in contrast to the directional tracking described for the α-factor receptor Ste2 (Wang et al., 2019). To compare the behavior of Ste2 and polarity probes, we imaged mating in cells harboring both *STE2-sfGFP* and either *BEM1-tdTomato* or *SPA2-mCherry*, a formin regulator that colocalizes with indecisive polarity sites (Henderson et al., 2019). Cortical Ste2 behavior varied, but most cells developed Ste2 crescents that gradually became broader and more intense (***Fig. 2E***). Neither Ste2 nor polarity factors displayed obvious directional tracking, as shown in three examples chosen to illustrate the range of behaviors (***Fig. 2E***). In one cell, the polarity factors exhibited unidirectional movement from the initial polarity site to the final site (*top*), in a second, initial polarization occurred near the partner with minimal movement (*middle*), and in a third, polarity factors showed erratic movement over the cortex (*bottom*). Taken together, these observations suggest that polarity factors initially concentrate at a small site on the cortex, then exhibit indecisive rather than stereotyped behavior before commitment.

### Pheromone sensing and secretion are enriched at transient polarity sites

Pheromone receptors and G proteins are concentrated near stable polarity sites (Ayscough and Drubin, 1998; McClure et al., 2015; Suchkov et al., 2010), but it was unclear whether the transient polarity sites characteristic of the indecisive period would similarly concentrate these factors. We imaged strains harboring both *BEM1-tdTomato* and either *STE2-sfGFP* (receptor) or *GFP-STE4* (Gβ) in mating mixes, identified time points at which a Bem1-tdTomato site was present, and assessed by visual comparison whether the Ste2 or Ste4 signal was colocalized. Both the receptor and Gβ were sometimes colocalized with polarity factors (***Fig. 3A***). The frequency of detectable colocalization during the indecisive period varied between cells (***Fig. 3B***), but these data suggest that even transient polarity sites can create detectable enrichment of pheromone sensing proteins (***Fig. 3C***).

**Fig. 3.**
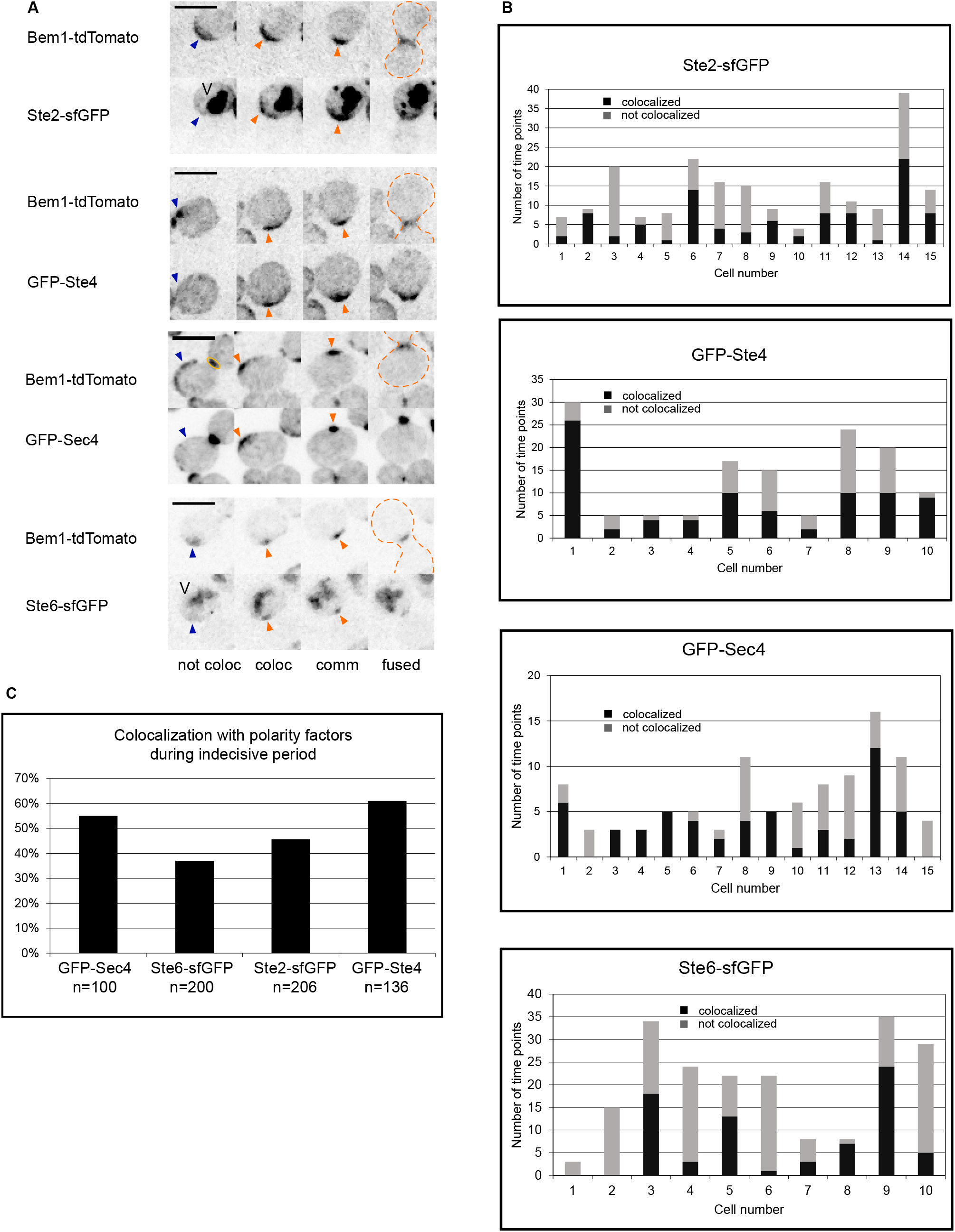
Localization of pheromone secretion, sensing, and signaling proteins during the indecisive period. Strains harboring *BEM1-tdTomato* and either the α-factor receptor *STE2-sfGFP* (DLY22243), Gβ subunit *GFP-STE4* (DLY23354), secretory vesicle marker *GFP-SEC4* (DLY13771), or a-factor transporter *STE6-sfGFP* (DLY22355) were imaged during mating. (A) Representative images show examples of the indicated probes during the indecisive period (left two images), in committed cells (comm) and just after fusion (fused). Internal signal in Ste2-sfGFP and Ste6-sfGFP strains is due to sfGFP accumulation in the vacuole (V) following Ste2/Ste6 degradation. For time points when cells had clusters of Bem1-tdTomato, the green probe was scored as either not colocalized (not coloc, blue arrowheads) or colocalized (coloc, orange arrowheads). Yellow oval: cytokinesis site. Orange dashed line: zygote. Scale bars: 5 μm. (B) The colocalization frequency during the indecisive phase, scored as illustrated in (A), varied from cell to cell. Different cells had indecisive periods of different durations, and only some of the time points showed clear clusters of Bem1 for scoring. Y-axis: number of time points scored per cell. (C) Overall colocalization frequency (% of time points during the indecisive phase that show colocalization of the indicated probe with the Bem1 signal). n, total number of time points scored.

We also examined whether pheromone secretion might occur preferentially at polarity sites. The two pheromones are secreted by different mechanisms: α-factor is delivered to the plasma membrane in secretory vesicles, and **a**-factor is secreted by a transporter, Ste6 (Michaelis and Barrowman, 2012). As with the pheromone sensing probes, GFP-Sec4 (a marker of secretory vesicles) and Ste6-sfGFP were often enriched at indecisive polarity sites (***Fig. 3A-C***).

It is unclear whether the degree of enrichment we observed would enable exploratory polarization. If much of the pheromone sensing and secretion is distributed around the cortex, then cells could also decode pheromone gradients using a global strategy, integrating the spatial distribution of pheromone over time and polarizing in the direction that provides the strongest signal. This model does not ascribe any role to the weak and transient polarity sites that are seen prior to commitment.

### Wildtype cells fail to commit to partners that lack polarity sites

To investigate the role of indecisive polarity sites, we asked how wildtype cells respond to partners that cannot polarize Cdc42. Because loss of Cdc42-GTP abrogates pheromone-responsive signaling (Simon et al., 1995; Zhao et al., 1995), we began by activating Cdc42 all over the cortex. To activate Cdc42, we overexpressed a membrane-targeted, constitutively-active guanine nucleotide exchange factor (GEF: *MT-GFP-CDC24^38A^*), a strategy previously shown to abrogate polarization (Kuo et al., 2014). To ensure that the mutants (mating type **a**) were arrested in G1, we added 10 μM α-factor. In arrested mutant cells, both Bem1-tdTomato and MT-GFP-Cdc24^38A^ were broadly distributed on the plasma membrane (***Fig. S1***).

In mating mixes with wildtype **a** cells, wildtype α cells (Bem1-tdTomato) formed polarity sites that varied in intensity, location, and number, typical of the indecisive period, before committing to a partner and fusing (***Fig. 4A, Movie 2***). However, when mixed with *MT-CDC24^38A^* mutants, wildtype α cells did not commit to the mutant partners, instead displaying extended indecisive behavior (***Fig. 4B, Movie 3***). Under these conditions, wildtype pairs mated at a high frequency during 2-h movies (mating efficiency ME = 0.62, n = 162). However, no wildtype cells were observed to mate with the unpolarized mutants (ME = 0, n = 151, p < 0.01), even when imaged for 3 h.

**Fig. 4.**
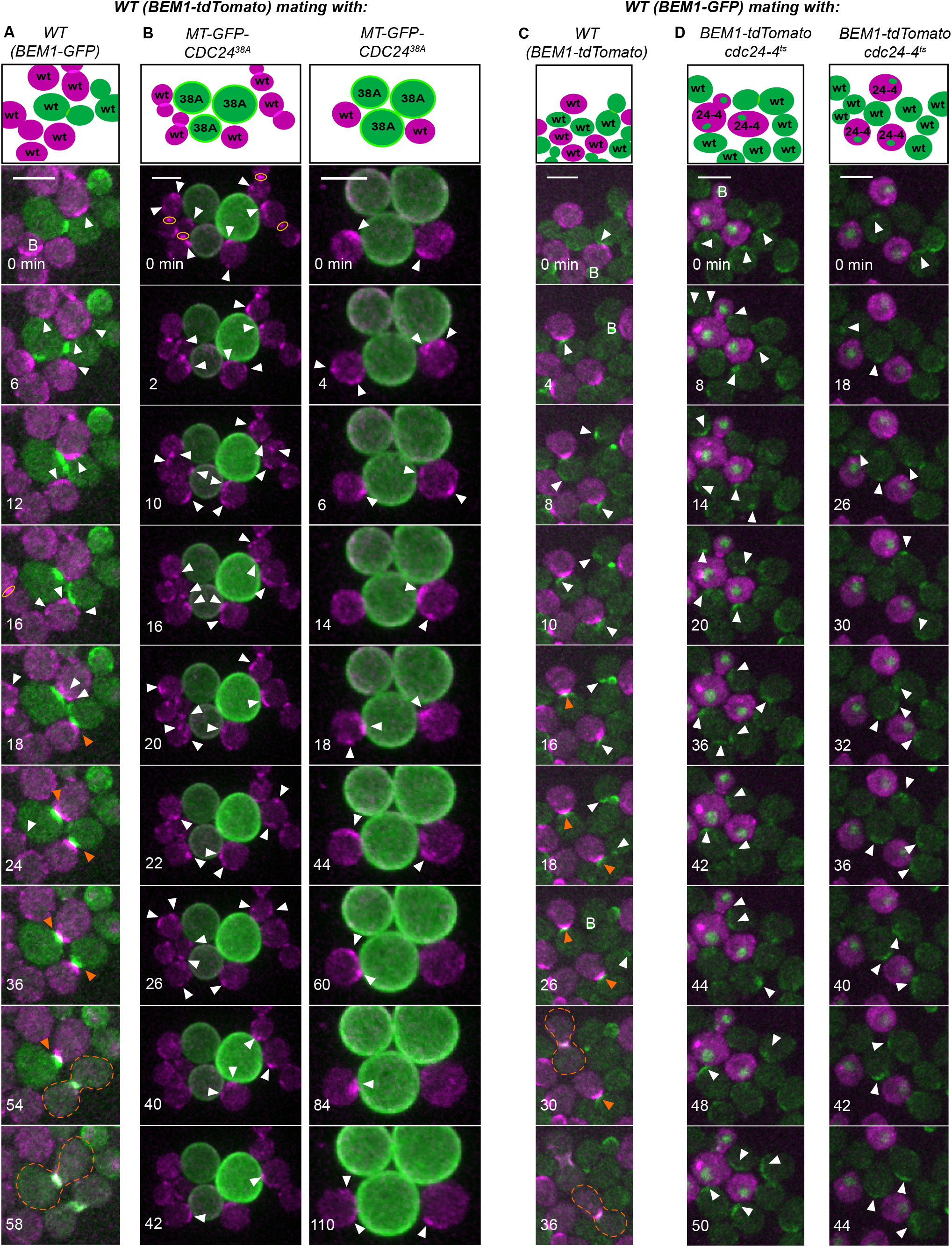
Wildtype cells do not commit to unpolarized partners. Selected time points from movies of mating mixes. Cartoons indicate cells in the selected montages at the start of the displayed imaging interval. B: bud. Yellow oval: mother-bud neck. White arrowhead: weak, mobile Bem1 cluster characteristic of indecisive cells, focusing on the magenta (A,B) or green (C,D) channel wildtype cells. Orange arrowheads: stably oriented Bem1 clusters characteristic of committed cells. Dashed outline: fused zygote. (A) MATα wildtype cells (*BEM1-tdTomato*, DLY12944) mixed with MAT**a**wildtype cells (*BEM1-GFP*, DLY9069), imaged at 30°C. (B) The same MATα wildtype strain mixed with MAT**a** cells harboring membrane-targeted, constitutively-active Cdc24 (*MT-GFP-CDC24^38A^,* DLY23351) that do not make polarity clusters and imaged at 30°C. Two montages are shown. (C) MATα wildtype cells (*BEM1-GFP*, DLY9070) mixed with MAT**a** wildtype cells (*BEM1-tdTomato*, DLY12943), imaged at 37°C. (D) The same MATα wildtype strain mixed with MAT**a** cells harboring *cdc24-4^ts^* (DLY23256, green nuclei indicate G1 cells), imaged at 35°C. Two montages are shown. Scale bars: 5 μm.

The failure of *MT-GFP-CDC24^38A^* mutants to communicate effectively might stem from excess Cdc42-GTP, rather than the lack of Cdc42 polarization. To address that possibility, we performed a similar experiment with strains lacking Cdc42-directed GEF activity. At the restrictive temperature, *cdc24-4^ts^* mutants fail to polarize and have predominantly inactive Cdc42-GDP (Adams et al., 1990; Atkins et al., 2013). However, *cdc24-4^ts^* mutants also fail to activate Ste20 and cannot respond to pheromone. We restored pheromone signaling by introducing the constitutively-active *ste20^ΔCRIB^*, which eliminates the need for Cdc42-GTP to respond to pheromone (Moskow et al., 2000). Again, we added 10 μM α-factor to ensure G1 arrest. At 35 or 37°C, wildtype cells (Bem1-GFP) displayed both indecisive and committed behavior in pairings with other wildtype cells (***Fig. 4C, Movie 4***), but when mixed with mutants, they again exhibited prolonged indecisive behavior (***Fig. 4D, Movie 5***). At 35°C, wildtype cells mated at high frequency with wildtype partners (ME = 0.80, n = 115), but did not mate with unpolarized mutants (ME = 0, n = 83, p < 0.01).

To quantify the behavior of the wildtype partners in these mixes, we developed an unbiased scoring method to distinguish indecisive and committed behavior based on the Bem1 probe. We calculated the correlation between the spatial distribution of pixel intensities in a cell between consecutive time points. If the polarity sites are mobile, the correlation is low, but if the cells commit, the correlation is high (***Fig. 5A***). In wildtype mating mixes, this “spatial autocorrelation” generally increased at commitment. 50 mating pairs were scored to select a commitment threshold that minimized false positives and false negatives (***Fig. S2A***). Because polarity sites were brighter and less variable at 30°C than 35°C, we chose a different threshold for each condition (***Fig. S2B***).

**Fig. 5.**
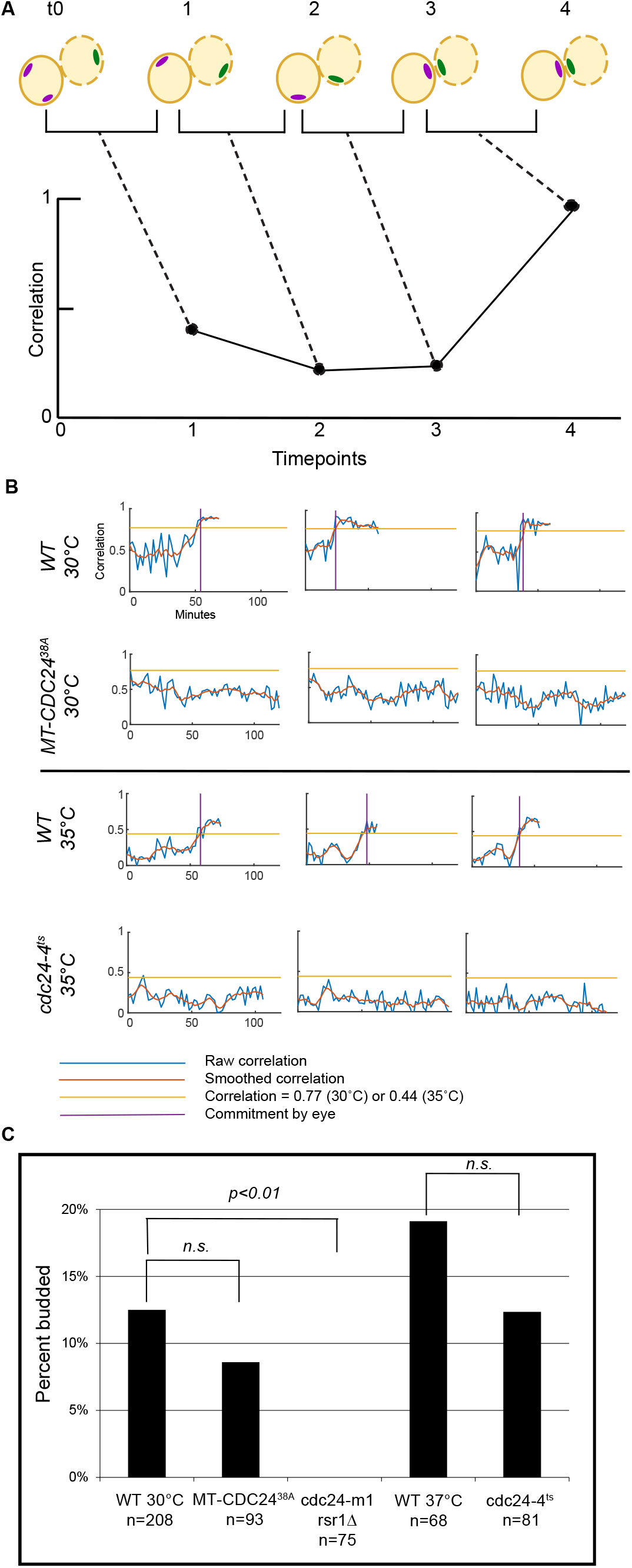
Scoring commitment and cell cycle arrest in mating mixes. (A) Cartoon illustrating spatial autocorrelation algorithm to score commitment. The spatial distribution of Bem1 pixel intensities in a cell of interest (magenta clusters) are compared at consecutive time points to yield a normalized correlation measure between 0 (no correlation) and 1 (perfect correlation). Strong and stably oriented polarity sites characteristic of committed cells (3,4) yield a high correlation while weaker, mobile polarity sites characteristic of indecisive cells (0,1,2) yield a low correlation. (B) Example spatial autocorrelation traces from wildtype cells mixed with either wildtype partners or the indicated non-clustering mutants as in ***Fig. 4***. Horizontal yellow line: threshold autocorrelation used to call commitment. Purple vertical line: commitment time as scored visually. Wildtype cells attempting to mate with unpolarized mutants did not reach the threshold, even after 100 min. (C) Wildtype cells adjacent to G1-phase cells of opposite mating type sometimes return to the cell cycle and form a bud. The percent of wildtype cells that budded was determined from the mating mixes with the indicated partners (genotypes as in ***Fig. 4*** and ***6***).

Whereas 48/50 cells mixed with wildtype partners developed spatial autocorrelation above threshold at 30°C, 0/20 cells mixed with *MT-CDC24^38A^* partners did so, despite prolonged imaging (***Fig. S3***). Similarly, 42/50 cells mixed with wildtype partners crossed the threshold at 35°C, but only 5/20 cells mixed with *cdc24-4^ts^* partners did so (***Fig. S4***). We conclude that wildtype cells do not commit to *MT-CDC24^38A^* or *cdc24-4^ts^* mutants as they do to wildtype partners (p < 0.01 for both). Examples of spatial autocorrelation traces are shown in ***Fig. 5B***.

Because unpolarized mutants might secrete less pheromone than wildtype cells, we assessed the capacity of the mutants to induce cell cycle arrest in wildtype partners in mating mixes, calculating the percentage of cells that budded during attempted mating. Similar percentages of wildtype cells budded when attempting to mate with either wildtype cells or unpolarized mutants (*MT-CDC24^38A^*, p = 0.29; *cdc24-4^ts^*, p = 0.17), demonstrating that the mutants secrete sufficient pheromone to trigger and maintain cell cycle arrest (***Fig. 5C***). We conclude that indecisive polarity sites are critical to communicate a cell’s location to its partner, and that without such communication, a wildtype cell does not commit.

### Wildtype cells fail to commit to constitutively indecisive partners

The exploratory polarization hypothesis posits that cells commit when they sense concentrated pheromone released from an immediately apposed partner polarity site. If consistently high local signaling is required to sustain commitment, then a wildtype cell would be unable to commit to a partner that was unable to stabilize the position of its own polarity site, even if that partner had mobile polarity sites.

Polarity sites can be stabilized by either of two parallel pathways, each of which recruits Cdc24 to the cortex (Dyer et al., 2013). One pathway depends on binding of the scaffold Far1 to Cdc24, which is impaired in the *cdc24-m1* mutant (Butty et al., 1998; Nern and Arkowitz, 1998; Nern and Arkowitz, 1999; Valtz et al., 1995). The other depends on the Ras-family GTPase Rsr1 (Bender and Pringle, 1989; Chant and Herskowitz, 1991). When treated with concentrated pheromone, mutants lacking both of these pathways (*cdc24-m1 rsr1Δ*) exhibit constitutively mobile polarity sites (Dyer et al., 2013; Nern and Arkowitz, 2000), and we expected that they would similarly exhibit constitutively mobile sites in a mating mix.

When mixed with wildtype α cells, *cdc24-m1 rsr1Δ* **a** cells formed constitutively mobile polarity sites, reminiscent of indecisive period behavior (***Fig. 6A, Movie 6***). Notably, their wildtype partners (Bem1-tdTomato) exhibited similar behavior, arresting in G1 and forming transient polarity sites but failing to commit. We asked if this phenotype was mating type-specific by repeating this experiment with wildtype **a** cells, and found that these also failed to commit to α partners with constitutively mobile polarity sites (***Fig. 6B,C***). Consistent with earlier work (Nern and Arkowitz, 1999), mating mixes between wildtype and *cdc24-m1 rsr1Δ* strains showed poor mating efficiency (ME = 0.01, n = 189, p < 0.01). As with the unpolarized mutants, 0/20 wildtype cells randomly selected for spatial autocorrelation analysis committed to *cdc24-m1 rsr1Δ* partners (***Fig. 6D***). Furthermore, no budding events were observed among wildtype cells that were adjacent to potential mating partners (n = 75), confirming that the pheromone signal released by *cdc24-m1 rsr1Δ* strains was sufficent to sustain arrest in wildtype potential partners (***Fig. 5C***). We conclude that effective communication with a partner requires that the partner possess indecisive polarity sites and is capable of stabilizing them.

**Fig. 6.**
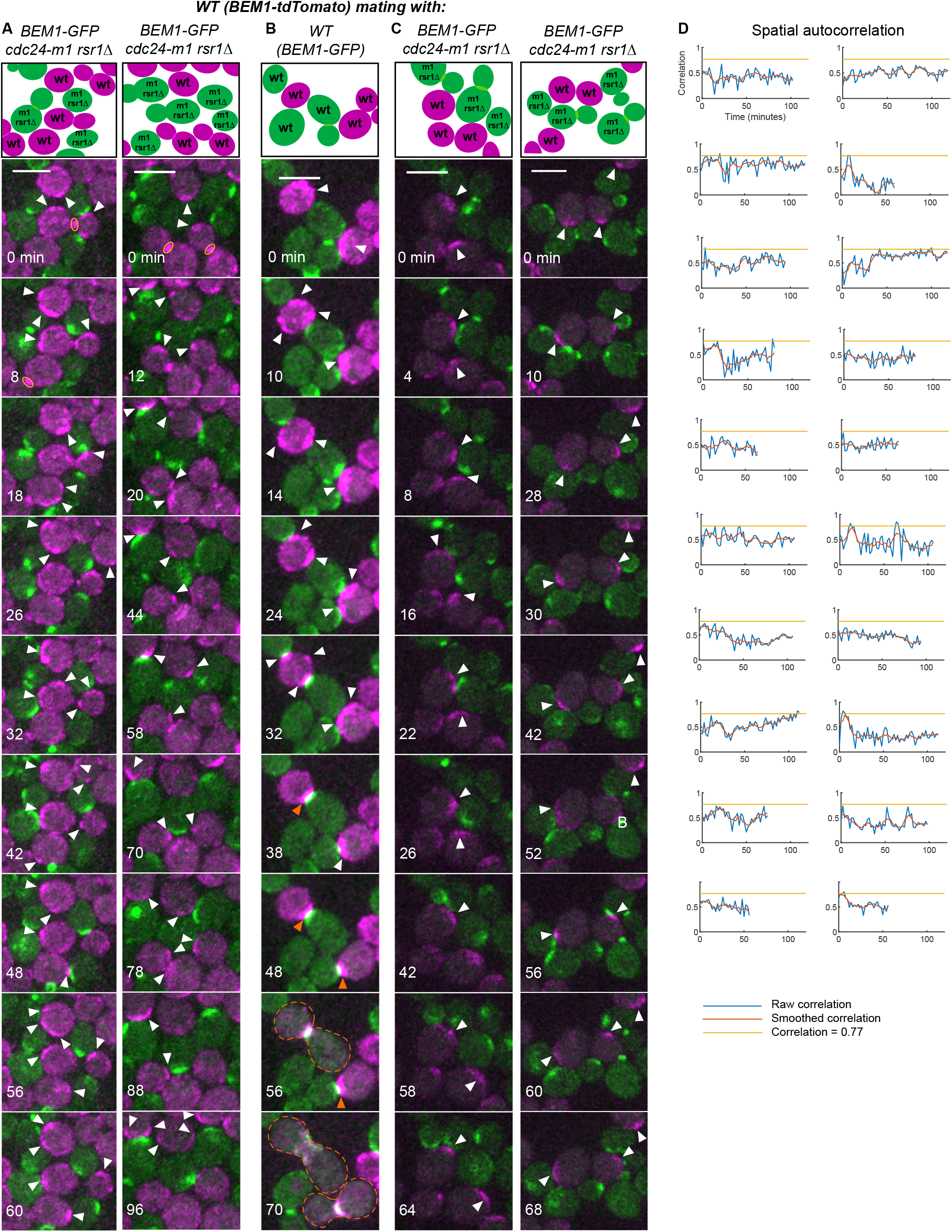
Wildtype cells do not commit to mutants with constitutively mobile polarity sites. (A-C) Selected time points from movies of mating mixes. Yellow oval: mother-bud neck. White arrowhead: weak mobile Bem1 cluster characteristic of indecisive cells, focusing on the magenta (A) or green (B,C) channel wildtype cells. Orange arrowheads: stably oriented Bem1 clusters characteristic of committed cells. Dashed outline: fused zygote. (A) MATα wildtype cells (*BEM1-tdTomato*, DLY12944) were mixed with MAT**a** mutants that form constitutively mobile polarity clusters (*cdc24-m1 rsr1Δ BEM1-GFP*, DLY22797) and imaged at 30°C. (B) MATα wildtype cells (*BEM1-GFP*, DLY9070) were mixed with MAT**a** wildtype cells (*BEM1-tdTomato*, DLY12943). Control mating mix in which mating type and fluorophore are switched relative to ***Fig. 4A***. (C) MAT**a** wildtype cells (*BEM1-tdTomato*, DLY12943) were mixed with MATα mutants that form constitutively mobile polarity clusters (*cdc24-m1 rsr1Δ BEM1-GFP*, DLY23612). (D) Spatial autocorrelation traces of representative wildtype cells mixed with *cdc24-m1 rsr1Δ* cells from (A).

### Wildtype cells can commit to partners that are stably oriented toward them

Previous studies indicated that mating can still occur, albeit at reduced frequency, in conditions where the **a**-type mating partner is “confused”—that is, unable to locate partner cells—due to the addition of saturating levels of α-factor to the medium (Dorer et al., 1997; Dorer et al., 1995). Saturating α-factor causes cell cycle arrest and stable polarization, but it is unclear how such cells mate. We speculated that the low frequency of successful mating in these conditions might reflect fortuitous instances in which the “confused” cell happens to orient directly toward an α partner, and that during its own indecisive phase, the α cell finds and commits to the confused cell. Alternatively, the presence of saturating α-factor may allow a “unilateral” mating in which only one partner needs to orient properly.

We imaged wildtype cells in a mating mix with 10 μM α-factor. **a** cells polarized stably and grew in a single direction which usually did not point to an α partner. The α cells next to such “misoriented” **a** cells exhibited prolonged indecisive behavior and did not commit or mate (***Fig. 7, Movie 7***). In the rarer instances in which an **a** cell polarized toward an α partner, some α cells polarized toward the **a** cell’s polarity site and mated (***Fig. 8A, Movie 8***). Thus, cells can mate with partners that stably orient in the correct direction, even if that orientation develops by chance and not through a search process. This accounts for all of the mating events we observed (n = 14). Curiously, cells were able to mate even if the polarity site of the non-confused partner was less stable than in typical pairings, as reflected in the fact that the spatial autocorrelation metric did not reach the commitment threshold (***Fig. 8C***). Cells that did not mate (regardless of orientation) never reached the commitment threshold (***Fig. 7B, 8C***). We did not observe any fusion events without preceding coorientation, suggesting that only if the two partners’ polarity sites are oriented toward each other do the cells commit and mate. Interestingly, we also observed instances where pairs that appeared to be properly oriented failed to mate under “confusion” conditions (n = 36) (***Fig. 8B, Movie 9***). The basis for this behavior remains unknown.

**Fig. 7.**
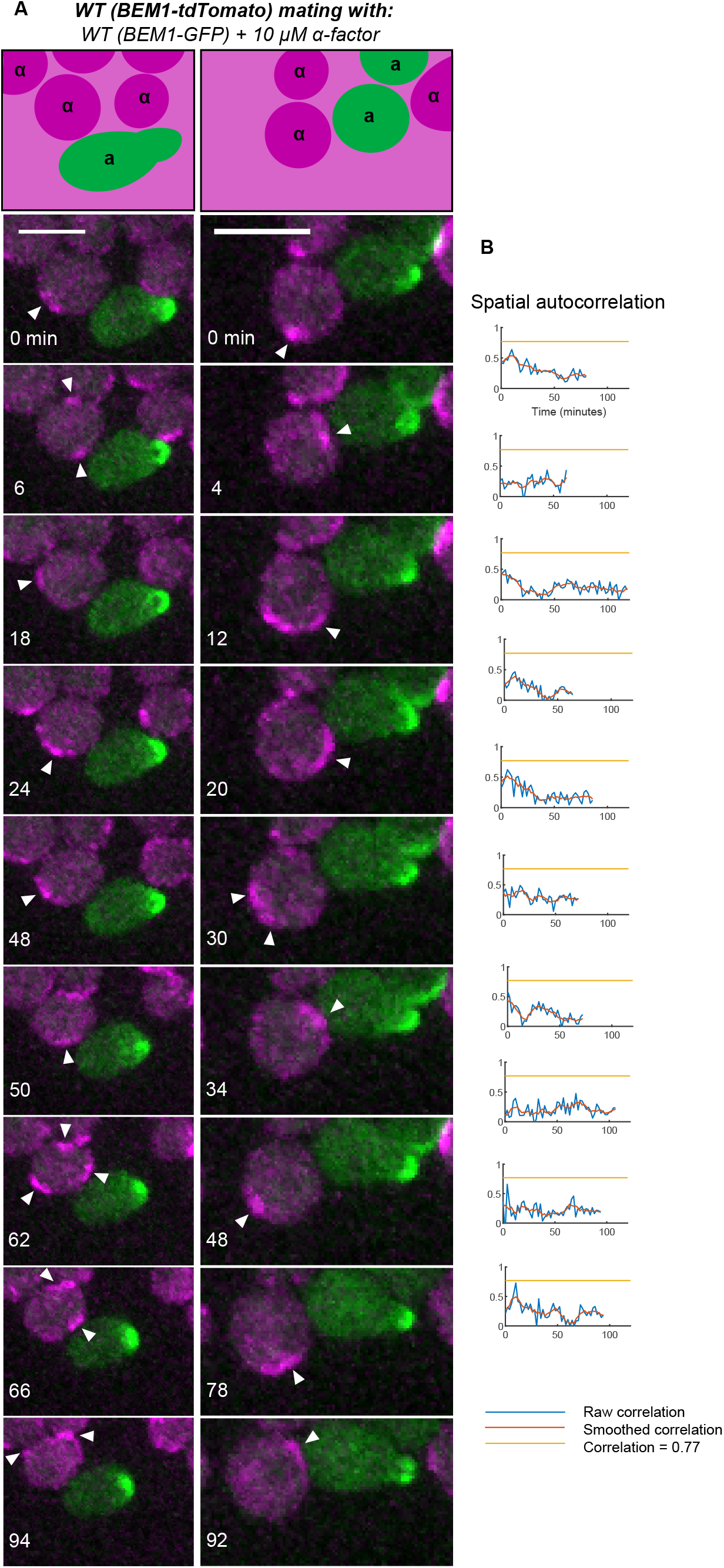
Wildtype cells do not commit to mutants with stable but misoriented polarity sites. (A) Selected time points from movies of mating mixes. White arrowhead:weak mobile Bem1 cluster characteristic of indecisive cells, focusing on the magenta channel wildtype cells. In the presence of 10 μM α-factor, MAT**a** cells (*BEM1-GFP*, DLY9069) polarize stably and form mating projections. When these projections are oriented away from the MATα wildtype potential partner (*BEM1-tdTomato*, DLY12944), the partner forms polarity sites that remain indecisive. (B) Representative spatial autocorrelation traces of MATα cells from (A).

**Fig. 8.**
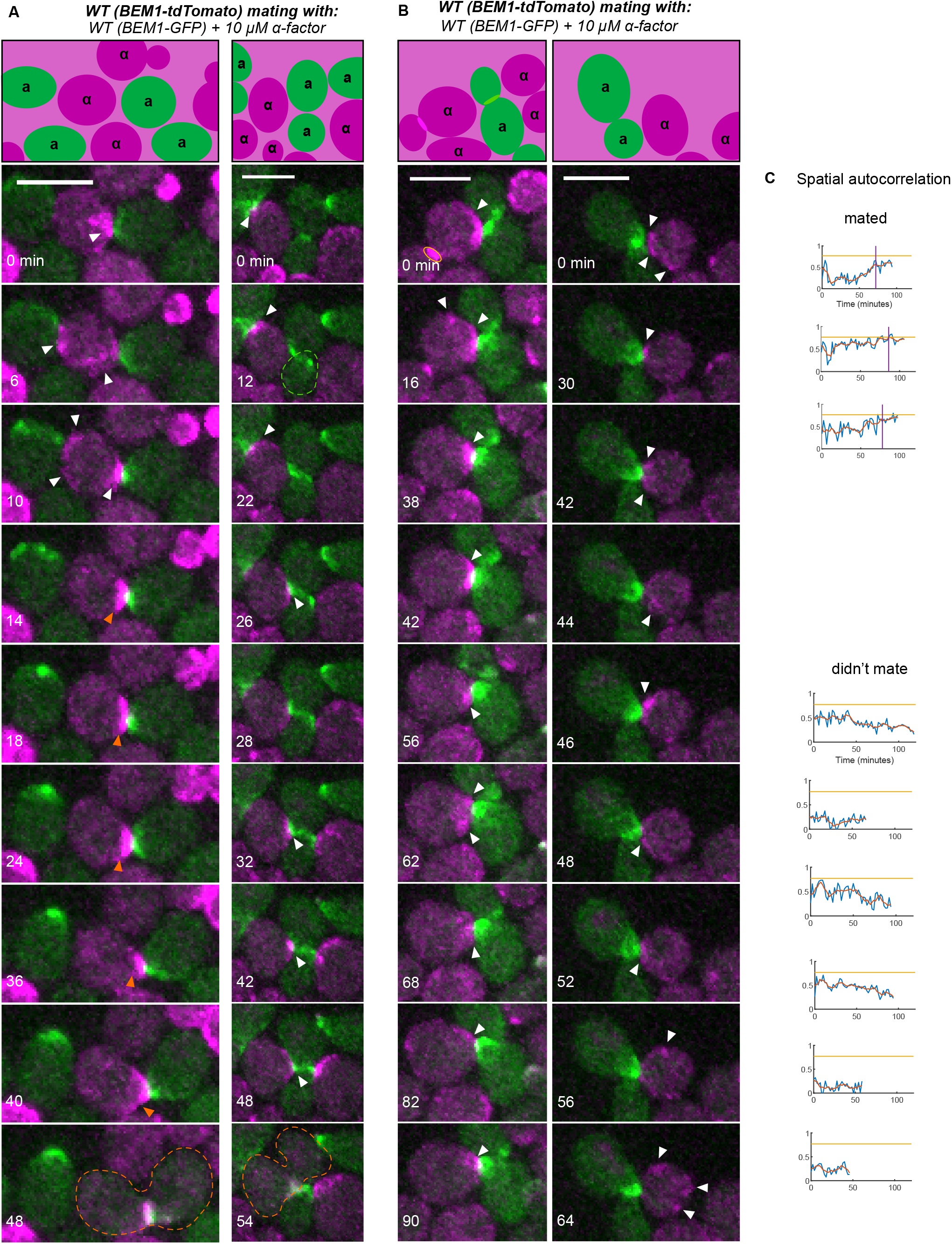
Default mating requires fortuitous correct orientation by the “confused” partner. (A,B) Selected time points from movies of the same mating mixes as in **Fig. 7**. (A) Examples of successful mating. (B) Examples in which mating fails despite apparently correct orientation by the confused partner. (C) Representative spatial autocorrelation traces of MATα cells that did (top three) or did not (bottom six) mate.

### Simulating the pheromone landscape experienced by mating cells

Our findings indicate that for yeast cells to commit to a mating partner, pheromone secretion must be oriented from each partner toward the other. Unpolarized or misdirected secretion did not trigger commitment. To understand this, we first estimated the pheromone concentration that might be detected by a partner next to a cell secreting pheromone. An α cell arrested in G1 following exposure to **a**-factor secretes approximately 1400 molecules of α-factor per second (Rogers et al., 2012). Assuming that the pheromone profile reaches steady state, a cell that secretes pheromone in an unpolarized manner would generate a local pheromone concentration of only 0.5 nM (Materials and Methods). However, if the same amount of pheromone were secreted in a focused manner from a small zone, then the local concentration could exceed 5 nM, comparable to the receptor K_D_ (Jenness and Spatrick, 1986). As stable polarization requires a higher pheromone concentration than cell cycle arrest (Hegemann et al., 2015; McClure et al., 2015; Moore, 1983; Paliwal et al., 2007), this calculation suggests that cells adjacent to unpolarized partners may fail to commit simply because they do not sense a sufficiently high concentration of pheromone.

To better understand how a “local sensing” cell that detects pheromone at a zone surrounding the polarity site would respond to an adjacent partner cell, we simulated an arrangement with two spheres: a pheromone emitter and a pheromone receiver. The spheres were 250 nm apart (the minimal possible distance based on the combined thickness of two cell walls) to simulate cells that are touching (***Fig. 9A***). α-factor is secreted by exocytosis of vesicles, which fuse at a rate of ~ 0.83/s (Dyer et al., 2013). Thus, with an overall α-factor release rate of 1400/s (Rogers et al., 2012), the average number of pheromone molecules in a vesicle would be ~1680.

**Fig. 9.**
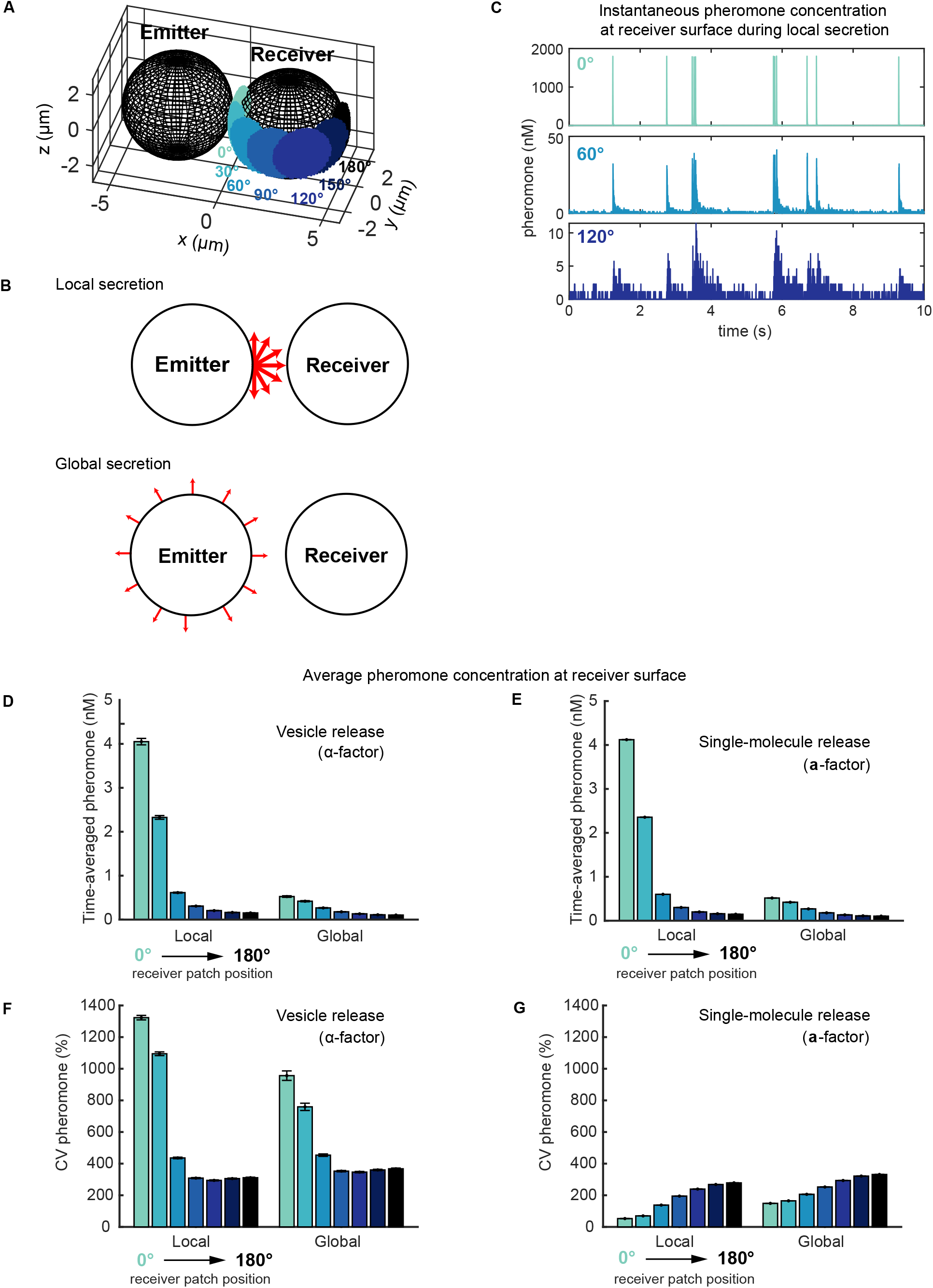
Simulations of the pheromone receiver’s landscape for two touching cells. (A) Model setup for emitter and receiver cells shown at scale. Seven patch positions on the receiver (0° to 180°, changing colors) were used to measure local pheromone concentrations. (B) Local versus global secretion. In local secretion, pheromone was released just at the emitter pole abutting the receiver. In global secretion, pheromone was released uniformly just at the emitter surface. (C) Instantaneous pheromone concentration at different positions (color) near receiver’s surface over time during local vesicle secretion. (D,E) Time-averaged pheromone concentration at different positions (color) on receiver’s surface for both vesicle and single-molecule release. (F,G) Coefficient of variation (CV) for (D,E). All bars show mean ± s.e.m., n = 300 realizations.

We simulated pheromone release in one of two patterns: global secretion, where each vesicle releases its pheromone at a random position on the surface of the emitter, or local secretion, where each vesicle releases pheromone at the pole that abuts the receiver (***Fig. 9B***). Following secretion, pheromone molecules were assumed to diffuse freely unless reflected from the surfaces of the two spheres. To simulate pheromone sensing in the vicinity of the polarity site, we designated a ~1.3 μm diameter patch at several locations (0° to 180°, changing colors, ***Fig. 9A***) on the receiver, and counted the number of molecules within 0.25 μm of the patch surface. Pheromone concentrations calculated in this manner fluctuated dramatically as vesicles were released (***Fig. 9C***).

Pheromone-receptor unbinding is slow (k_off_ = 0.01-0.001/s) (Bajaj et al., 2004; Jenness et al., 1983; Raths et al., 1988; Yi et al., 2003), suggesting that receptors would average the local pheromone concentration rather than responding rapidly to a transient spike. Temporal averaging of pheromone concentrations in different patches on the receiver indicated that the concentration sensed in the patch facing the emitter was ~8-fold higher for simulations with polarized secretion compared to those with global secretion, consistent with the estimates discussed above (***Fig. 9D***). Thus, if a threshold concentration must be detected to promote commitment, then cells that secrete pheromone in a polarized manner would be much more likely to cross that threshold. Furthermore, the pheromone concentrations sensed at different locations declined much more steeply with distance from the emitter in the simulations with polarized secretion (***Fig. 9D***). Thus, when secretion and sensing both occur at polarity sites, the concentration sensed by a cell would depend on the relative positions of the two cells’ polarity sites, as posited by the exploratory sensing hypothesis.

Unlike α-factor, **a**-factor is exported by the transporter Ste6 (Michaelis and Barrowman, 2012), so that **a**-factor release may occur one molecule at a time (Michaelis and Barrowman, 2012), rather than in vesicular packets. We repeated the simulations discussed above, assuming the same overall production rate but releasing one molecule of pheromone at a time. While the variability in pheromone concentration was greatly reduced (***Fig. 9F***), the average pheromone concentrations sensed at different locations remained the same (***Fig. 9E***).

## DISCUSSION

### Exploratory polarization underlies partner selection in yeast mating

Our findings indicate that the transient polarity sites formed during the indecisive period are critical for subsequent commitment to a mating partner. Proteins involved in pheromone sensing, secretion, and signaling were all enriched at these sites, suggesting that they are preferred sites for both pheromone secretion and sensing. Wildtype cells in mating mixes did not commit to partners that lacked polarity sites, partners with constitutively mobile polarity sites, or partners with stable but misoriented polarity sites. These results are fully consistent with the exploratory polarization hypothesis (***Fig. 1D***), in which transient polarity sites mediate communication between mating partners.

Previous findings indicated that the appearance of a strong, stable polarity site, which we call “commitment,” results from detection of a high concentration of pheromone (Hegemann et al., 2015; Henderson et al., 2019; McClure et al., 2015; Moore, 1983). Our findings suggest that pheromone levels sufficient to trigger commitment are only achieved when a partner’s polarity site is directly apposed to that in the receiving cell. Simulations based on reported pheromone production rates confirm that local pheromone secretion would expose a well-oriented polarity site to much higher pheromone levels (4 nM) than those attainable by a cell secreting pheromone uniformly around its surface (0.5 nM). We conclude that yeast cells normally commit to a partner in response the concentrated pheromone signal that accompanies coorientation of the two cells’ polarity sites.

An open question concerns the mechanism whereby the two partner cells’ polarity sites “find each other” to become cooriented. One could imagine that polarity sites form, move, and disappear stochastically until coorientation promotes stable commitment, as proposed for “speed dating” in *S. pombe* (Bendezu and Martin, 2013; Merlini et al., 2016). Alternatively, polarity sites may be guided toward each other by pheromone gradients. Our simulations indicate that when pheromone is secreted locally, the mating partner would experience a very steep gradient in pheromone concentration, potentially guiding the movement or formation of polarity sites.

A recent study suggested that initial polarity sites move gradually and directionally toward the partner cell in *S. cerevisiae* (Wang et al., 2019). In our mating conditions, the movement was more chaotic, without obvious unidirectional tracking in most cells. Nevertheless, it remains possible that the movement is influenced by the pheromone landscape in a manner that accelerates coorientation. Such movement could occur via local sensing of pheromone gradients near the polarity site or global sensing of pheromone all over the surface.

### Re-evaluating the pheromone landscape of mating cells

The observation that yeast cells are able to orient polarization toward artificial pheromone sources generated by micropipets (Nern and Arkowitz, 1998; Segall, 1993; Valtz et al., 1995) or microfluidic devices (Brett et al., 2012; Dyer et al., 2013; Hao et al., 2008; Hegemann et al., 2015; Jin et al., 2011; Kelley et al., 2015; Lee et al., 2012; Moore et al., 2008; Moore et al., 2013; Paliwal et al., 2007; Vasen et al., 2020) has focused attention on the mechanism whereby cells decode a stable gradient of pheromone. Although yeast cells are clearly capable of polarizing growth toward an exogenous pheromone source, wildtype cells failed to polarize growth toward partners that were secreting pheromone uniformly around their surface. As such cells would be expected to set up a stable pheromone gradient similar to that from a micropipet, why did their partners not commit?

Experiments that analyze polarization in artificial pheromone gradients generally focus on cells that remain arrested in G1 for prolonged periods (4-10 h). Cells that are further from the pheromone source arrest only transiently and then resume budding, and these cells are omitted from the analysis of directional polarization in the gradient. However, we suggest that this transiently arrested population may be the most relevant to the behavior of mating cells. In our wildtype by wildtype matings, we found that 12-19% of cells went on to bud during a 2-h observation window (***Fig. 5C***). Note that this analysis excludes cells that were not directly adjacent to (touching) potential G1-phase mating partners. Thus, cells that do not mate are unlikely to remain arrested in G1 for many hours under these circumstances. The simplest explanation for mating failures with unpolarized partners is that yeast cells simply do not secrete enough pheromone to recreate the kinds of gradients produced by microfluidics devices.

Unlike experimental settings with unidirectional gradients, yeast cells in physiological mating scenarios must often discriminate between two or more similarly-distant pheromone sources (McClure et al., 2018; Taxis et al., 2005). Under those circumstances, stable pheromone gradients would seem unlikely, and the findings presented in this and other recent studies (Bendezu and Martin, 2013; Dyer et al., 2013; Henderson et al., 2019; Merlini et al., 2016; Wang et al., 2019) suggest that mating cells operate in the context of a fluctuating pheromone landscape quite unlike the stable gradients studied thus far. Fluctuations occur on several timescales. First, vesicular release of α-factor would generate dramatic spikes in pheromone concentration, because each vesicle contains very concentrated (~ 4 mM) α-factor. However, with an estimated α-factor diffusion constant of 150 μm^2^/s, each spike would dissipate to low nM levels very rapidly (< 0.1 s, ***Fig. S5A***), well before the next spike, generating large second-to-second fluctuations. Second, the movement of the polarity sites during the indecisive phase means that the source of pheromone would relocate on a minute timescale, shifting the local gradients. Third, on a several-minute timescale, the mating or budding of nearby cells would remove them as pheromone sources in the local environment. Thus, physiological pheromone gradients are likely to be transitory, at least until the partners commit to each other. We suggest that the exploratory polarization strategy provides a framework for understanding how yeast cells are able to locate partners and mate successfully in such a dynamic pheromone landscape.

### Advantages of exploratory polarization

The exploratory polarization strategy supported by our findings, like the related speed dating strategy proposed for *S. pombe* (Bendezu and Martin, 2013; Martin, 2019; Merlini et al., 2016), provides an elegant solution to the problem of choosing a partner from among two or more similarly-distant candidates. Classical spatial sensing paradigms that integrate spatial information to extract a single “up-gradient” direction are poorly suited to this task, as the presence of two or more nearby chemoattractant sources may create a weak or even non-existent net gradient. The task of picking just one of the potential partners is accomplished by the coincidence-detection feature of exploratory polarization: stabilization of the polarity site only occurs when the partners’ polarity sites happen to align (***Fig. 1D***). By including this temporal aspect in the partner search process, the cells can avoid the potential paralysis that could ensue from access to two or more equally attractive partners. We note that this partner selection task occurs not only in mating, but also more broadly in multicellular contexts that involve focal cell-cell junctions like synapses.

A potential problem with exploratory polarization stems from the observation that during the indecisive phase, cells frequently developed two or more transient polarity sites. In principle, then, a cell could end up with two polarity sites each oriented toward a different partner, leading to double mating. How cells avoid this problem poses an interesting question for future investigation.

## Acknowledgements

We thank Stefano Di Talia, Masayuki Onishi, and Amy Gladfelter, as well as members of the Lew lab for stimulating conversations and comments on the manuscript. M.R.C-C. was a Howard Hughes Medical Institute Gilliam Fellow and received a Graduate Diversity Enrichment Program Award from the Burroughs Wellcome Fund. This work was funded by NIH/NIGMS grants R35GM127145 to T.C.E. and R01GM103870 and R35GM122488 to D.J.L.

## MATERIALS AND METHODS

**Table S1.**
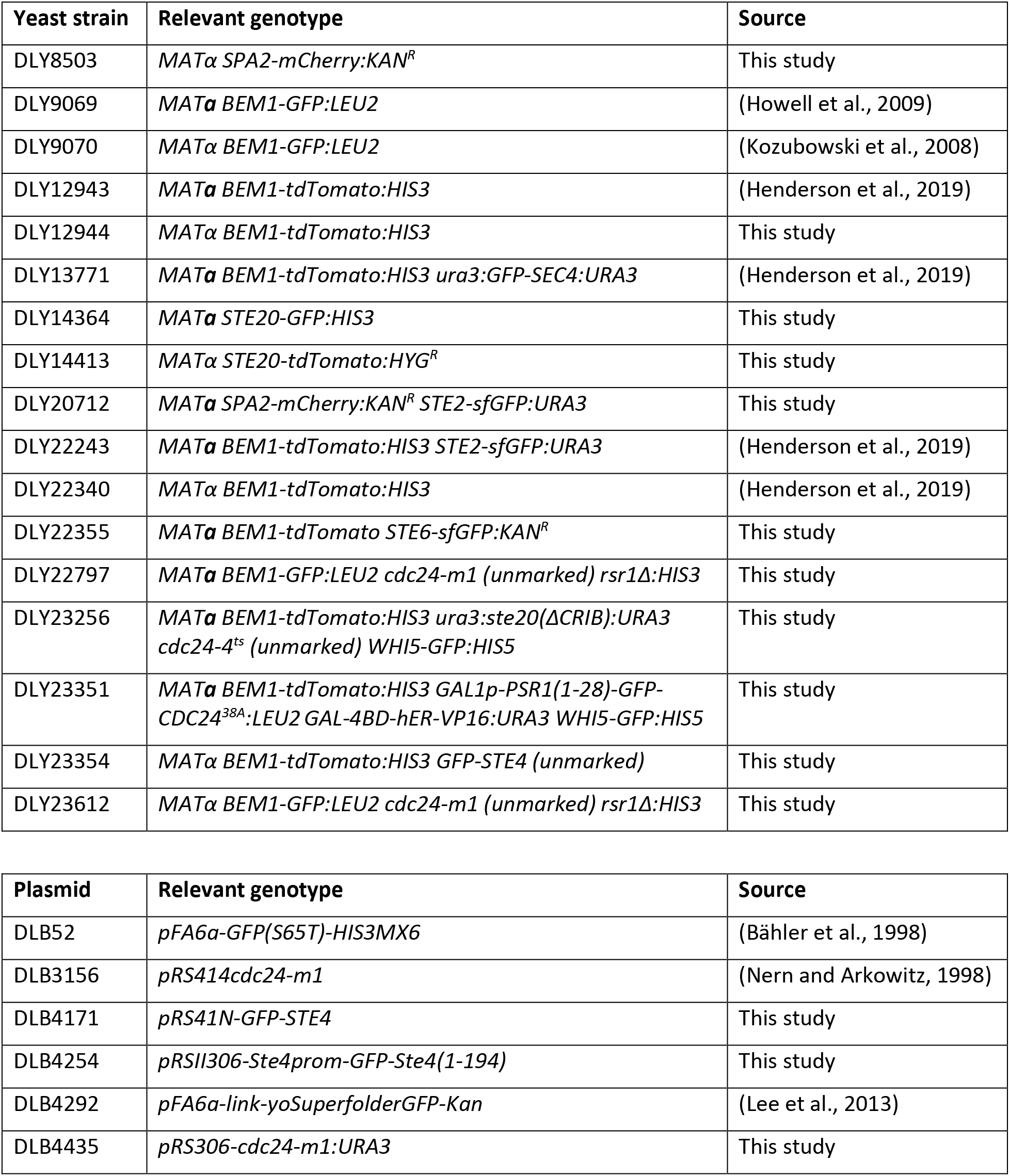
Yeast and plasmid strains and genotypes.

### Yeast strains and plasmids

Strains were constructed using standard molecular biology techniques. Yeast strains used in this study (***Table S1***) were generated in the YEF473 background (*his3-Δ200 leu2-Δ1 lys2-801 trp1-Δ63 ura3-52*) (Bi and Pringle, 1996). The following alleles were previously described: *BEM1-GFP:LEU2* (Kozubowski et al., 2008), *BEM1-tdTomato:HIS3* (Howell et al., 2012), *SPA2-mCherry:KAN^R^* (Howell et al., 2009), *GFP-SEC4:URA3* (Chen et al., 2012), *STE20-GFP:HIS3* and *ura3:ste20(ΔCRIB):URA3* (Moran et al., 2019), *STE2-sfGFP:URA3* (Henderson et al., 2019), *rsr1Δ:HIS3* (Schenkman et al., 2002), and *GAL1p-PSR1(1-28)-GFP-CDC24^38A^:LEU2* (Woods et al., 2015).

*WHI5-GFP:S.p.HIS5* (Doncic et al., 2011) and *STE6-sfGFP:KAN^R^* were constructed using methods described previously (Longtine et al., 1998) with DLB52 (pFA6a-GFP(S65T)-S.p.HIS5MX6; Addgene plasmid #41598) and DLB4292 (pFA6a-link-yoSuperfolderGFP-Kan; Addgene plasmid #44901) as templates.

*GFP-STE4* was constructed in the YEF background by a pop-in, pop-out strategy. First, *GFP-STE4* was PCR-amplified from a strain derived from RDY126 (Suchkov et al., 2010). This fragment was inserted into *pRS41N* (Taxis and Knop, 2006) using ApaI and NotI, producing DLB4171 (*pRS41N-GFP-STE4*). DLB4171 was digested with HindIII and ApaI to release a fragment containing the *STE4* promoter, GFP, and bp 1-194 of the ORF. This fragment was inserted into DLB212 (*pRSII306:* a *URA3-*marked integrating plasmid) to produce DLB4254 (*pRSII306-Ste4prom-GFP-Ste4(1-194)*). DLB4254 was partially digested with PstI to target integration to the *STE4* locus of a diploid from the YEF background. Haploid segregants containing *STE4(1-194)*, *URA3* marker, and GFP-*STE4* at the *STE4* locus were plated on medium containing 5-fluoroorotic acid to select for colonies in which recombination occurred between the promoter of the *STE4* fragment and the promoter of *GFP-STE4*, removing the *URA3* marker and leaving *GFP-STE4* as a precise replacement of the endogenous *STE4*.

To introduce the *cdc24-4* allele into the YEF473 strain background, we first deleted one copy of *CDC24* in a diploid strain using the *HIS3* marker. A centromeric *URA3*-marked plasmid carrying wildtype *CDC24* was transformed into the strain, and following sporulation and tetrad dissection a haploid *cdc24::HIS3* strain carrying the plasmid was selected. *cdc24-4* was amplified by PCR from a strain derived from JPT19-H01 (Sloat et al., 1981), and used to replace the *cdc24::HIS3* allele by homologous recombination, followed by selection on medium containing 5-fluoroorotic acid to obtain colonies without the plasmid.

The *cdc24-m1* allele was amplified by PCR from *pRS414-cdc24-m1* (Nern and Arkowitz, 1998) and cloned into pRS306 to produce DLB4435 (*pRS306-cdc24-m1*). DLB4435 was digested with BspEI to target integration at the *CDC24* promoter, yielding a locus where the *URA3* gene is inserted between the *cdc24-m1* allele and wildtype *CDC24*. Haploid MAT**a** segregants were grown on medium containing 5-fluoroorotic acid to select for recombination between *cdc24-m1* and *CDC24*. Recombinants containing *cdc24-m1* were identified by phenotyping (morphology when treated with pheromone) and confirmed by sequencing.

### Live-cell microscopy

Cells were grown in complete synthetic medium (CSM; MP Biomedicals, LLC, Solon, OH, USA) with 2% dextrose (Macron, Center Valley, PA, USA) overnight at 30°C to mid-log phase (10^6^ – 10^7^ cells/ml). Cultures of opposite mating type strains were mixed to obtain a 1:1 cell ratio, centrifuged to concentrate the cells, and mounted on CSM-dextrose slabs solidified with 2% agarose (Hoefer, Holliston, MA, USA) and sealed with petroleum jelly. For pheromone “confusion” experiments, cells were imaged on a slab containing 10 μM α-factor (Genway Biotech, San Diego, CA, USA). *GAL1p-MT-CDC24^38A^* expression was induced in dextrose medium by adding β-estradiol (Sigma-Aldrich, St. Louis, MO, USA) to the medium to a final concentration of 20 nM, incubating for 2 h, and imaging on a slab containing 20 nM β-estradiol (the strains contain an artificial transcription factor, *GAL-4BD-hER-VP16*, that induces the *GAL1* promoter in response to β-estradiol). For *cdc24-4^ts^* strains, cells were grown overnight at 24°C and shifted to 37°C for 2 h before imaging. For cells that were arrested in G1 (*MT-CDC24^38A^* or *cdc24-4^ts^*), α-factor was added to a final concentration of 10 μM, and cells were incubated for 2 h before imaging on a slab containing 10 μM α-factor. Imaging was performed in a temperature-controlled chamber at 30°C, 35°C or 37°C as indicated.

Images were acquired with an Andor Revolution XD spinning disk confocal microscope (Andor Technology, Concord, MA, USA) with a CSU-X1 5,000-rpm confocal scanner unit (Yokogawa, Tokyo, Japan) and a UPLSAPO 100×/1.4 oil-immersion objective (Olympus, Tokyo, Japan), controlled by MetaMorph software (Molecular Devices, San Jose, CA, USA). Images were captured by an iXon3 897 EM-CCD camera with 1.2x auxiliary magnification (Andor Technology).

Images were acquired in z-stacks (15 0.47-μm steps) at 2-min intervals. Laser power varied by experiment but was set to levels that produced bright signals with minimal bleaching during the movie: 8-15% (488 nm) and 10-15% (561 nm) of maximal output. EM gain was 200, and exposure time was 250 ms. Images were denoised with the ImageJ Hybrid 3D Median Filter plug-in (2007), created by Christopher Philip Mauer and Vytas Bindokas. Images are maximum projections except for *MT-CDC24^38A^* medial plane image, as indicated. Scaling of images was always matched for experimental and relevant control conditions.

### Ste20 initial orientation analysis

From mating movies, we identified all cells in G1. We segmented each cell into four quadrants (one proximal, one distal, and two medial; ***Fig. 2A***), visually assigned the orientation of each cell’s initial Ste20 polarity site to a quadrant, and calculated the fraction of initial sites that formed in each quadrant. For the subpopulation analyses, cells were considered either “adjacent” to a potential partner in G1 or “isolated” (i.e. not adjacent).

### Colocalization analysis

From mating movies, we identified all cells in G1 that were adjacent to at least one potential G1-phase partner. For individual cells, we identified time points in which a clear localized Bem1 signal was present and scored the Ste2, Ste4, Sec4, or Ste6 signal as “colocalized” if the protein of interest was concentrated in the same area as Bem1 or “not colocalized” otherwise. For each pair of proteins, all time points were summed to calculate an overall colocalization frequency.

### Mating efficiency

Mating efficiencies were calculated from mating movies. We tracked all cells in G1 that were adjacent to at least one potential G1-phase partner (i.e. “available” to mate) during a 2-h movie and scored them as “mated” or “not mated.” Cells whose potential partners mated with another cell and cells that were available to mate only during the last 20 min were excluded from the sample. For each movie, three stage positions (i.e. technical replicates) were counted and summed to determine the fraction mated out of all available to mate (“mating efficiency”).

### Budding index

Budding indices were calculated from mating movies. Using the appearance of Bem1-GFP or Bem1-tdTomato at the neck as a marker of G1, we first calculated the duration of G1 for all wildtype α-cells that mated (mutants were **a**-cells) and determined the third quartile of that value (70 min). We then tracked all cells in G1 that were adjacent to at least one potential G1-phase partner (i.e. “available” to mate) for 70 min, noting whether they budded, mated, or remained arrested. The fraction of cells that budded was the budding index.

### Spatial autocorrelation analysis

The image processing toolbox in MATLAB 2019a was used to develop a custom tool to track individual cells during the mating period and determine commitment to a partner in an unbiased way. From mating movies, we identified wildtype α-cells (*BEM1-GFP* or *BEM1-tdTomato*) that were available to mate with **a**-cell partners. We circumscribed the wildtype cells at several time points throughout the movie, beginning at G1 and ending either at the time point preceding fusion (for cells that mated) or the end of the movie (for cells that did not mate). Using linear interpolation, the outlines were deformed over time to accommodate changes in cell morphology and position. In this manner, we obtained cell outlines between the marked time points, enabling continuous tracking of each cell during the mating period.

The spatial array of intracellular Bem1 signal was extracted at each time point, and the correlation between arrays at adjacent time points was calculated (“spatial autocorrelation”), using the following formula:

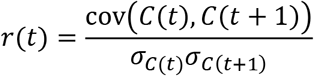

where *r* is Pearson’s coefficient, cov is covariance, *C* is an array containing indexed fluorescence data, *t* is time point, and *σ*_*C*_ is the standard deviation of the array *C*. The two arrays *C*(*t*) and *C*(*t* + 1) were obtained by using a union of the outlines at time points *t* and *t* + 1 to ensure that spatial overlap was continuous.

During the indecisive period, *r* is relatively low, but at commitment, *r* is relatively high. To determine a threshold of *r* that indicates commitment, we performed a sweep through different threshold values for a set of cells for which an experienced rater had already judged the time of commitment. A threshold value was selected to minimize discrepancies (either early or late judgments) between the automated and the human rater. The code used for this analysis can be found at: https://github.com/DebrajGhose/Exploratory-polarization-yeast

### Statistics

Except for spatial autocorrelation analysis, statistical G-tests of goodness-of-fit were used to compare groups (McDonald, 2014).

### Calculation of pheromone concentrations expected for global and local secreting cells

To gain insight into the types of gradients expected from global and local secreting cells, we considered the gradient generated by a spherical emitter centered at the origin (Rappaport and Barkai, 2012). For this case, the pheromone gradient can be found by solving Laplace’s equation using spherical coordinates. The r-coordinate satisfies the equation:

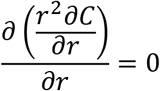

The boundary conditions are constant flux density J (number of molecules released per unit time per unit area) at the surface of the sphere (r = R)

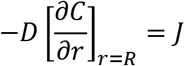

and the concentration vanishes as *r* → ∞. With these boundary conditions, the concentration takes the following form:

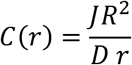

In terms of the total secretion rate S, the above expression becomes

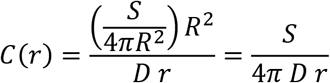

If we assume a secretion rate of 1400 molecules/s and diffusion coefficient for pheromone molecules of 150 μm^2^/s, then the concentration at the surface of cell of radius 2.5 μm is C(2.5) = 0.3 molecules/μm^3^ = 0.5 nM. If we assume the localized emitter has a radius of 0.25 μm, then the concentration at the surface of the emitter is 10 times higher C(0.25) = 5 nM. These results are consistent with the emitter alone results shown in ***Fig. S5B***.

### Simulations of pheromone landscape for two touching cells

Particle-based simulations of pheromone emission and diffusion were conducted using the Smoldyn software (v2.58) on Linux systems (2.50 GHz and 2.30 GHz Intel processors, Longleaf cluster at UNC Chapel Hill, NC, USA (Andrews, 2017; Andrews and Bray, 2004). Code is available at https://github.com/mikepab/exploratory-polarization-yeast. Pheromone molecules were modeled as Brownian point particles with diffusion coefficient D = 150 μm^2^/s, and were removed at a spherical absorbing boundary 50 μm from the origin. A mating pair was modeled as a two spheres, a receiver and emitter, centered at (±(2.5+0.25/2) μm, 0 μm, 0 μm) with radius 2.5 μm. The system was first equilibrated for 5 seconds, after which coordinates were recorded every 0.1 ms timestep for 10 seconds. For each condition, n = 300 realizations were conducted.

Vesicle emission events were simulated by repeated use of the Smoldyn command cmd @ t pointsource pheromone n x y z. First, t specifies the time of a single emission event; intervals between each emission were exponentially distributed:

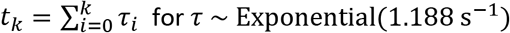

pheromone is a molecular species defined in the Smoldyn script, n is the number of molecules released per vesicle (n = 1663), and x y z are spatial coordinates of the vesicle event. In the local case, (x = −0.25/2+0.001 μm, y = z = 0 μm). In the global case, the spatial coordinates were obtained by uniform random sampling on a sphere centered at (−(2.5+0.25/2) μm, 0 μm, 0 μm) with radius 2.5001 μm.

Single-molecule emission events were handled using the Smoldyn reaction surface= and reaction_production commands, with a release rate per timestep of 1/7.1429 (yielding 1400 molecules per second at 0.1 ms timesteps). In the local case, the releasing surface was a sphere centered at x = −0.25/2+0.001 μm, y = z = 0 μm with a radius of 0.0005 μm. In the global case, the releasing surface was a sphere centered at (-(2.5+0.25/2), 0, 0) μm with radius 2.5001 μm.

To validate the simulation setup, we set up simulations comparable to the analytic solution described above. The receiver was removed and pheromone profiles were measured as a function of distance to the emitter. The simulations were in good agreement with the analytic solution (***Fig. S5B***).

### Analysis of particle-based pheromone simulations

Receiver-centered molecular coordinates were filtered to only include pheromone within 0.25 μm of the receiver surface. Then, a 3D angle between each molecule 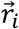 and a reference vector 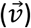 was calculated:

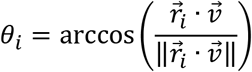

The reference vector defines the patch under consideration (Fig 9A). For 0°, the closest patch to the emitter 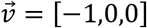. For the other patches, we rotate [−1,0,0] by the desired angle *θ*_*rot*_.

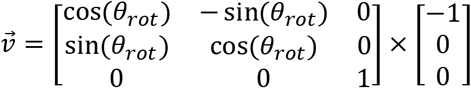

To count molecules in each patch, we summed the number of points within 0° ≤ *θ*_*i*_ ≤ 30°. The volume of each patch is patch is 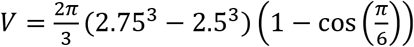, which was used to convert molecules to nanomolar. Finally, a time-averaged pheromone concentration and coefficient of variation (CV) were calculated for each patch in each simulation, allowing us to compute a mean and standard error across simulations (***Fig. 9D-G***).

## SUPPLEMENTARY FIGURE LEGENDS

**Fig. S1.**
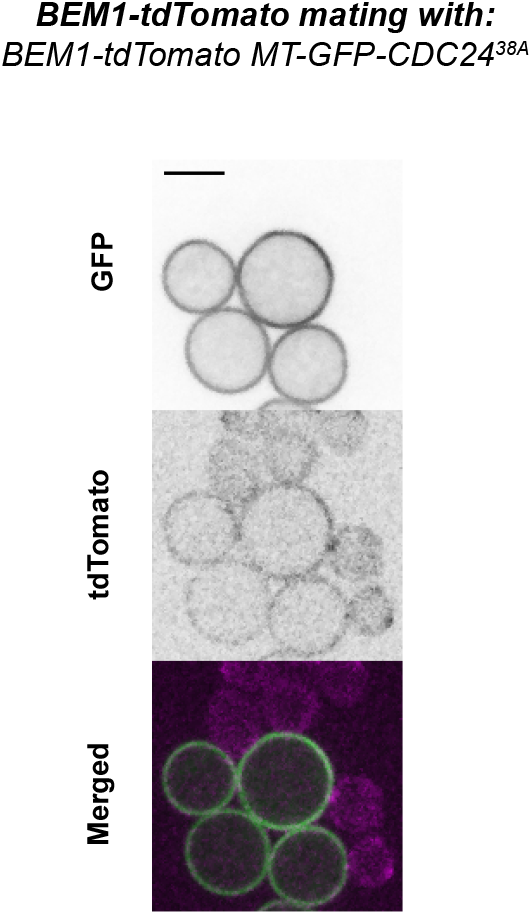
Overexpression of membrane-targeted Cdc24 blocks polarization. Medial plane confocal images of cells induced to express membrane-targeted, phospho-site mutant *GFP-CDC24^38A^* (*MT-GFP-CDC24^38A^ BEM1-tdTomato*, DLY23351) and mixed with wildtype cells (*BEM1-tdTomato,* DLY12944). Scale bar: 5 μm.

**Fig. S2.**
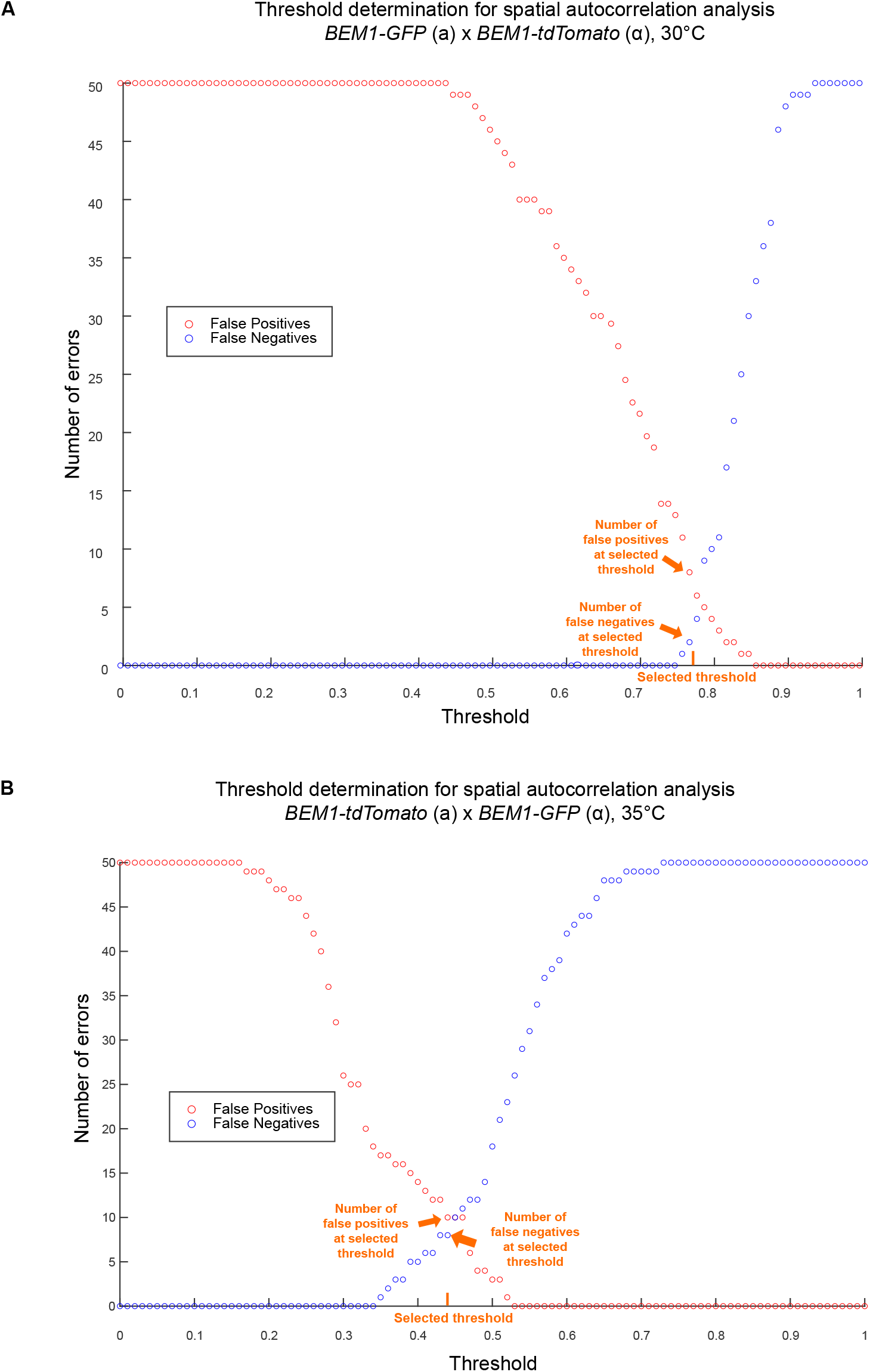
Threshold determination for spatial autocorrelation analyses. (A) The number of false negatives (in which the spatial autocorrelation trace did not cross the threshold but did commit as scored visually) and false positives (in which the spatial autocorrelation trace crossed the threshold > 4 min before commitment as scored visually) as a function of commitment threshold for wildtype x wildtype pairs at 30°C. A threshold of 0.77 was selected (orange tick). (B) Similar analysis for wildtype x wildtype pairs at 35°C. A threshold of 0.44 was selected (orange tick).

**Fig. S3.**
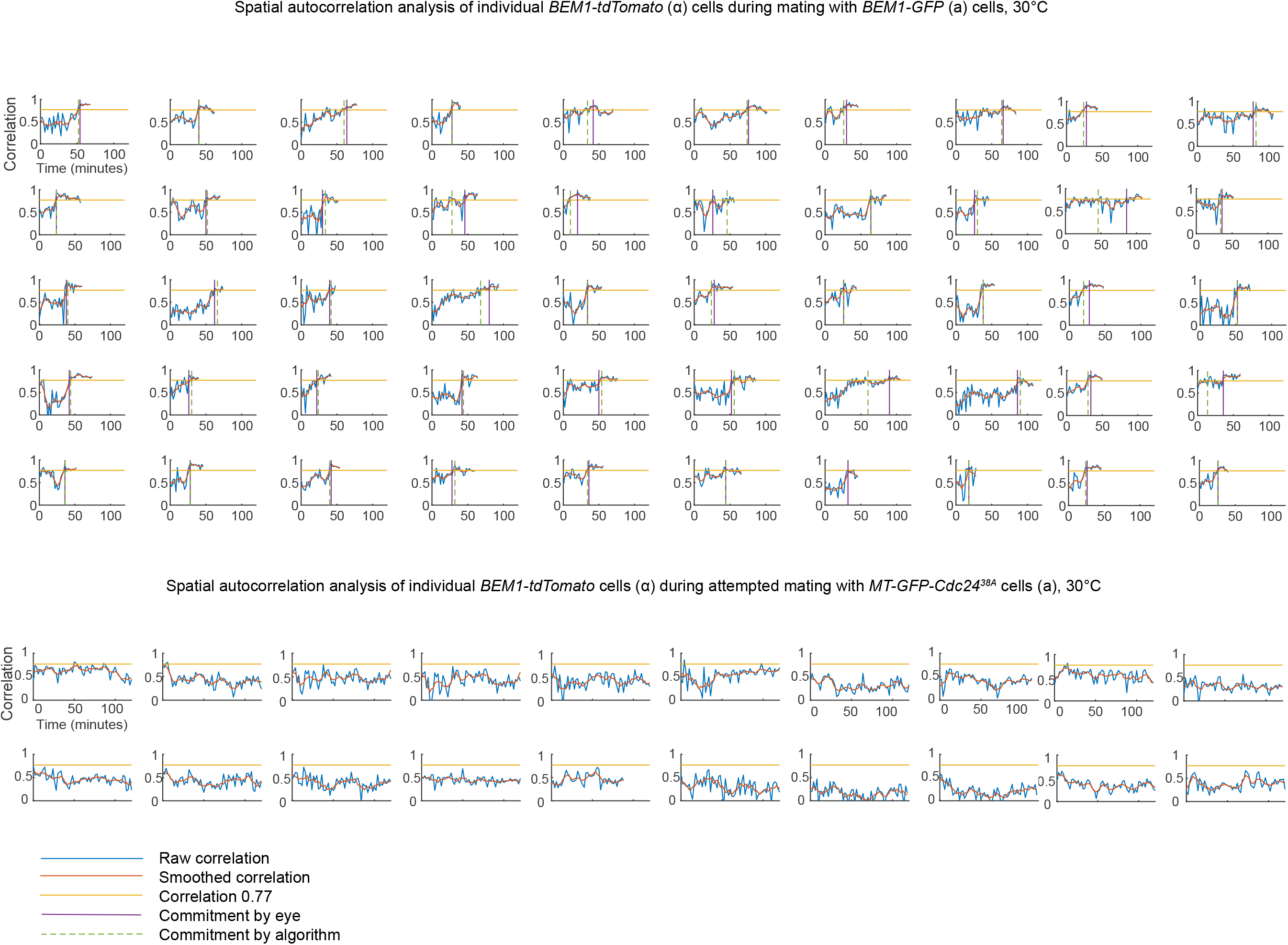
Spatial autocorrelation traces for cells at 30°C. (A) Wildtype by wildtype mixes as in Fig. 4A. Traces begin at the time of the cell’s entry into G1 and end at the time point preceding fusion. X-axis: Time (min). Y-axis: spatial autocorrelation (yellow line: commitment threshold). Commitment to a partner as determined visually (vertical purple line) or by crossing the threshold (dashed green line). (B) Similar analysis for wildtype cells mating with *MT-Cdc24^38A^* partners as in Figure 4B.

**Fig. S4.**
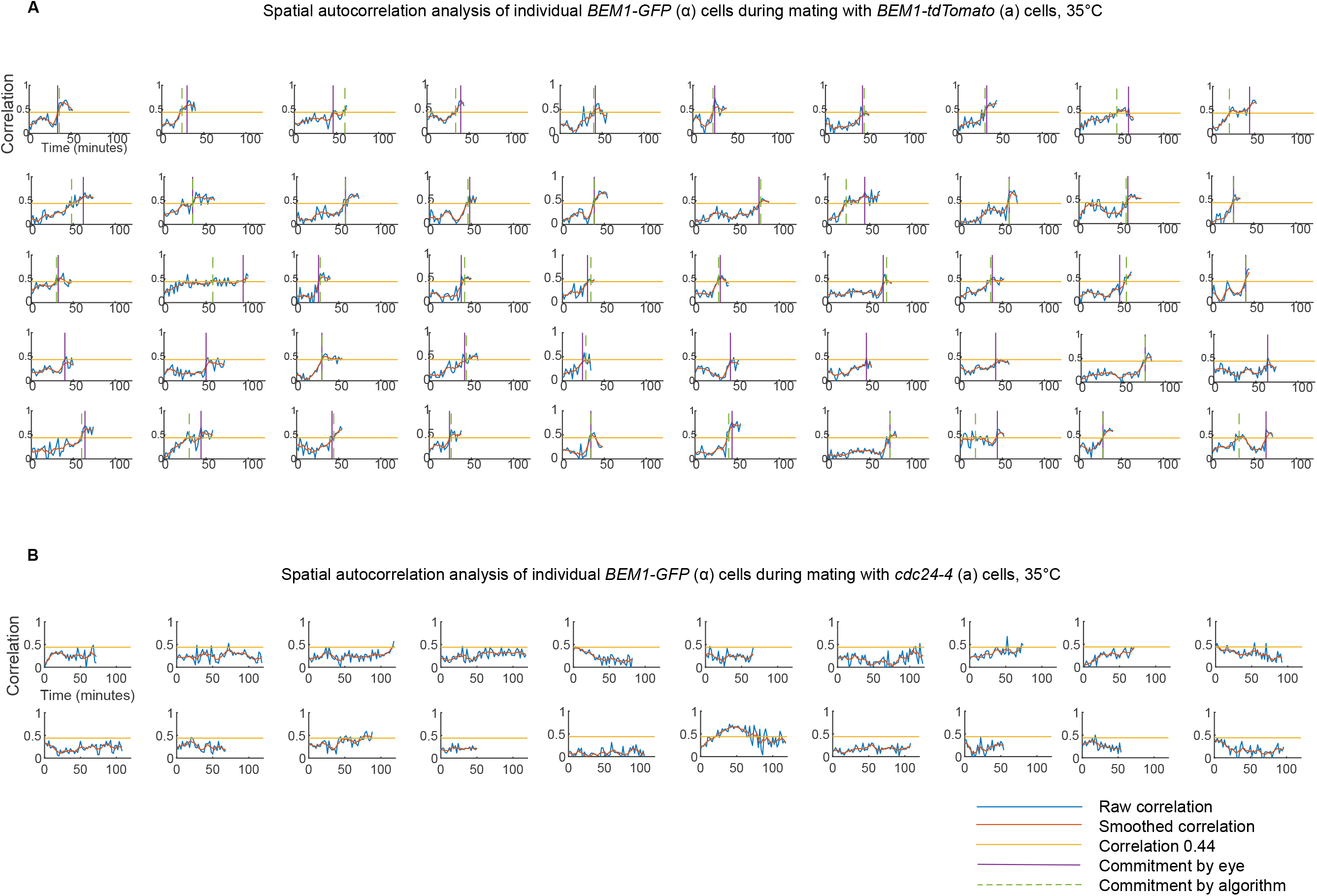
Spatial autocorrelation traces for cells at 35°C. (A) Wildtype by wildtype mixes. Traces begin at the time of the cell’s entry into G1 and end at the time point preceding fusion. X axis: Time (min). Y axis: spatial autocorrelation (yellow line: commitment threshold). Commitment to a partner as determined visually (vertical purple line) or by crossing the threshold (dashed green line). (B) Similar analysis for wildtype cells mating with *cdc24-4^ts^* partners as in Fig. 4D.

**Figure S5.**
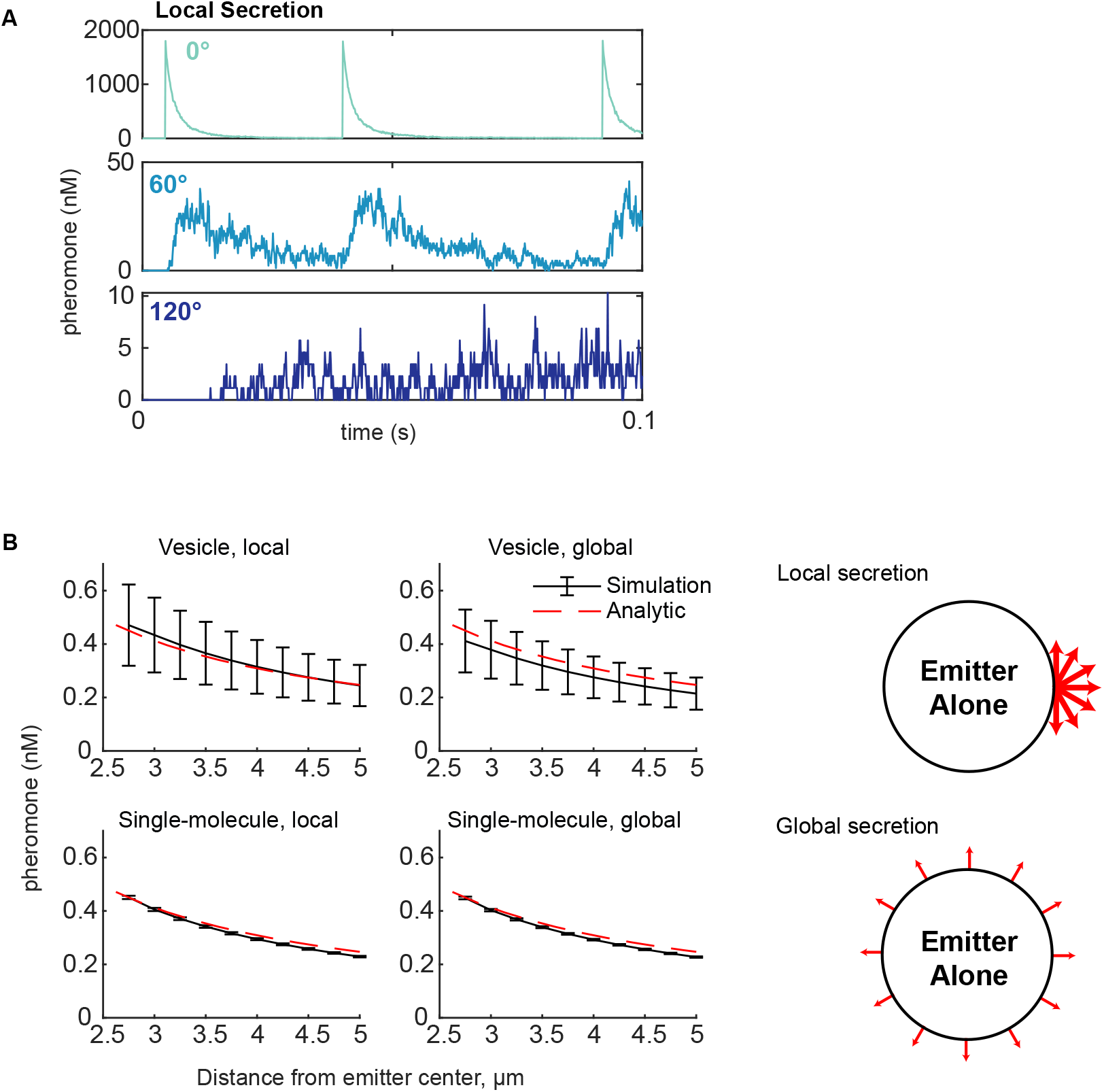
Validation of the pheromone simulations and additional detail. (A) Pheromone concentrations perceived at three different patches in a single simulation as in Fig. 9C, but zoomed in to show 0.1 second along the x-axis. (B) Simulations of the emitter alone, comparing concentrations in a spherical 250 nm shell (not a patch) at the indicated distance from the center of the emitter versus the steady-state analytic solution of the diffusion equation under equivalent conditions. Bars show mean ± s.d., n = 30.

## MOVIE LEGENDS

***Movie 1.*** Mating between cells harboring *STE20-GFP* (DLY14364) and *STE20-tdTomato* (DLY14413). Denoised maximum projections. Time in h:min. White arrowheads: selected polarity sites; orange arrowhead: commitment; dashed orange outline: zygote. Scale bar: 5 μm.

***Movie 2.*** Mating between cells harboring *BEM1-GFP* (DLY9069) and *BEM1-tdTomato* (DLY12944). Denoised maximum projections. Time in h:min. White arrowheads: selected polarity sites; orange arrowhead: commitment; dashed orange outline: zygote. Scale bar: 5 μm.

***Movie 3.*** Unsuccessful mating attempt between cells harboring *MT-GFP-CDC24^38A^ BEM1-tdTomato* (DLY23351) and *BEM1-tdTomato* (DLY12944). Denoised maximum projections. Time in h:min. White arrowheads: initial polarity sites. Scale bar: 5 μm.

***Movie 4.*** Mating between cells harboring *BEM1-tdTomato* (DLY12943) and *BEM1-GFP* (DLY9070). Denoised maximum projections. Time in h:min. White arrowheads: selected polarity sites; orange arrowhead: commitment; dashed orange outline: zygote. Scale bar: 5 μm.

***Movie 5.*** Unsuccessful mating attempt between cells harboring *cdc24-4^ts^* (DLY23256) and *BEM1-GFP* (DLY9070). Denoised maximum projections. Time in h:min. White arrowheads: initial polarity sites. Scale bar: 5 μm.

***Movie 6.*** Unsuccessful mating attempt between cells harboring *cdc24-m1 rsr1Δ* (DLY22797) and *BEM1-tdTomato* (DLY12944). Denoised maximum projections. Time in h:min. White arrowheads: initial polarity sites. Scale bar: 5 μm.

***Movie 7.*** Unsuccessful mating attempt between cells harboring *BEM1-GFP* (DLY9069, MAT**a**, “confused”) and *BEM1-tdTomato* (DLY12944, MATα, searching) plus 10 μM α-factor. Denoised maximum projections. Time in h:min. White arrowheads: initial polarity sites. Scale bar: 5 μm.

***Movie 8.*** Mating between cells harboring *BEM1-GFP* (DLY9069, MAT**a**, “confused”) and *BEM1-tdTomato* (DLY12944, MATα, searching) plus 10 μM α-factor. Denoised maximum projections. Time in h:min. White arrowheads: selected polarity sites; orange arrowhead: commitment; dashed orange outline: zygote. Scale bar: 5 μm.

***Movie 9.*** Unsuccessful mating attempt between cells harboring *BEM1-GFP* (DLY9069, MAT**a**, “confused”) and *BEM1-tdTomato* (DLY12944, MATα, searching) plus 10 μM α-factor. Denoised maximum projections. Time in h:min. White arrowheads: selected polarity sites. Scale bar: 5 μm.

